# Globetrotting geckos: Historical biogeography suggests an Indian origin and “Out-Of-India” dispersal for the cosmopolitan *Hemidactylus* geckos

**DOI:** 10.1101/2024.04.10.588862

**Authors:** Madhura Agashe, Aritra Biswas, K. Praveen Karanth

## Abstract

**Aim:** Following the breakup of Gondwana, dispersal events both into and out of India have influenced the biotic assembly of the surrounding landmasses. The cosmopolitan *Hemidactylus* presents an intriguing group to examine these instances, particularly given its sister relationship with the endemic *Dravidogecko* found in India’s Western Ghats. Despite earlier theories of Afro-Arabian or Southeast Asian origins for *Hemidactylus*, its deep divergence from *Dravidogecko* (∼57 million years ago) suggests a potential Indian origin, thus contradicting an “Into-India” scenario proposed for the current Indian tetrapod groups. We thus aim to resolve the origins and shed light on the inter-continental dispersals in *Hemidactylus* by reconstructing its biogeographic history.

**Location:** Worldwide

**Taxon:** *Hemidactylus* Geckos

**Methods:** We use six nuclear genes to reconstruct a reduced representation backbone phylogeny of *Hemidactylus* using likelihood, Bayesian, and coalescent based methods. We further assemble a timetree using a concatenated dataset of nuclear and mitochondrial markers from 132 of the 192 *Hemidactylus* species by constraining the topology with the backbone phylogeny and secondary calibrations. We use this chronogram to reconstruct ancestral geographic ranges using BioGeoBEARS, employing a time-stratified approach in conjunction with plate tectonics information to explicitly test four hypotheses regarding the origin of the genus — An Indian origin, a Southeast Asian origin, a Saharo-Arabian origin, or an Afrotropical origin.

**Results:** Our findings suggest that the ancestral lineage of *Hemidactylus* and *Dravidogecko* colonized the drifting Indian plate from Southeast Asia approximately 57 million years ago, eventually evolving into the two genera in the Indian subcontinent. Following this, *Hemidactylus* dispersed multiple times from India to Africa and Asia.

**Main Conclusion:** This study proposes an Indian origin for these widely distributed geckos, representing a rare instance of an “Out-of-India” dispersal scenario observed in a non-Gondwanan squamate group. Additionally, our research underscores the significance of incorporating sister taxa in biogeographic analyses to avoid misinterpretations of ancestral ranges.

## Introduction

The Indian subcontinent’s unique geological history, stemming from its Gondwanan origin and subsequent drift and collision with the Eurasian plate, has played a crucial role in facilitating the dispersal and diversification of different groups across landmasses (Karanth 2006). As the Indian Plate drifted northward, it created opportunities for faunal interchanges and dispersal of organisms between regions. It has been hypothesized that after the separation of India from Madagascar and Africa, it served as a “Biotic ferry” for the ancient Gondwanan groups that continued to evolve and diversify in India (Briggs 1989; 2003a; Bossuyt & Milinkovitch, 2001). During its course northward, dramatic geological and climatic changes due to the latitudinal shift of the landmass as well as the Deccan trap volcanic event likely wiped out most of these groups in India, including squamates (Mani 1974; Karanth 2006; 2021). With the collision of India and Eurasia, there was an exchange of fauna between the two landmasses, with surviving Gondwanan lineages dispersing out of India into Asia, and the Asian lineages dispersing into India (Datta-Roy & Karanth, 2009).

By analyzing molecular data, ancestral area reconstructions, and divergence times of various taxa, previous studies have attempted to unravel these intricate patterns of species dispersal and evolution, shedding light on the dynamic biogeography of India and Southeast Asia. Groups such as forest scorpions (Loria & Pendini, 2020), Ranidae frogs (Bossuyt & Milinkovitch, 2001), and Caecilians (Gower et al., 2002) have provided evidence supporting the “Out-Of-India” dispersal scenario.

On the other hand, the “Into-India” pattern has been shown for majority of the Indian taxa, including squamates such as Chameleons (Tolley et al., 2013), Draconine lizards (Grismer et al., 2016), and Varanids (Vidal et al., 2012). An extensive review carried out by Karanth (2021) using available time trees for the Indian tetrapods also suggests that the extant Indian squamates originated in Asia, dispersing into India following the collision of the Indian plate with Eurasia.

The gecko genus *Hemidactylus* Goldfuss (1820) forms an excellent model to test these two competing dispersal hypotheses since 1) The Indian subcontinent harbours almost one-third of the *Hemidactylus* diversity, 2) There are several endemic Indian species, 3) The genus currently has a wide distribution, and finally 4) These geckos are commonly known to undergo transoceanic dispersals.

*Hemidactylus* is a highly speciose and widespread genus of the family Gekkonidae, with 192 species currently recognized (Uetz et al.,2023). These geckos inhabit diverse habitats across all continents except Antarctica (Ceriaco et al.,2020). Additionally, they have established populations in major oceanic islands and archipelagos within tropical and subtropical regions (Vences et al., 2004). The extensive distribution of many *Hemidactylus* species has been attributed to their ability to undergo natural transoceanic dispersals (Kluge,1969). Further, species such as *H.mabouia*, *H.frenatus*, and *H.flaviviridis* are human commensals and have thus been inadvertently introduced in regions such as North America and Australia where they were naturally absent (Bauer et al., 2010, Agarwal et al., 2021). All members of this genus broadly show similarity in their toe morphology, however, there is considerable variation in external features such as body size, dorsal pholidosis, femoral pore presence and extent, and lamellar count (Carranza & Arnold, 2006). Due to their ubiquitous presence and unique characteristics, these geckos are excellent models for investigating various ecological and evolutionary hypotheses.

Recent research on *Hemidactylus* has focused primarily on taxonomy and systematics (e.g. Ceriaco et al., 2020; Smid et al., 2020; 2023; Lobon-Rovira et al., 2021; Adhikari et al., 2022; Das et al.,2022; Pal & Mirza, 2022; Khandekar et al., 2023; Narayanan et al., 2023), with fewer studies on biogeography (Lajmi et al., 2019; Murdoch, 2022), diversification (Lajmi & Karanth, 2020; Luzete et al., 2022), invasive potential and dispersal ecology (Agarwal et al., 2021; Rato et al., 2021; Lamb et al.,2021; Lapwong et al., 2021), and ectoparasite biology (Borroto-Páez et al., 2020; Diaz et al., 2020). Although many of these studies involve phylogenetic reconstructions, they are mostly localized. This has resulted in the resolution of species positions within each clade but has resulted in a gap in our understanding of the inter-clade relationships in this group.

Earlier investigations that attempted to resolve the phylogenetic placement of various clades have generated conflicting results with weak support for inter-clade relationships (Carranza & Arnold, 2006; Bauer et al.,2008, Bansal & Karanth, 2010; Bauer et al.,2010; see Fig 1). Additionally, these studies have been constrained by limited taxon sampling. Furthermore, there has been a flux of new species descriptions and taxonomic reassessments in the last decade, which needs to be included to better establish the phylogeny of this genus.

**Figure 1.**
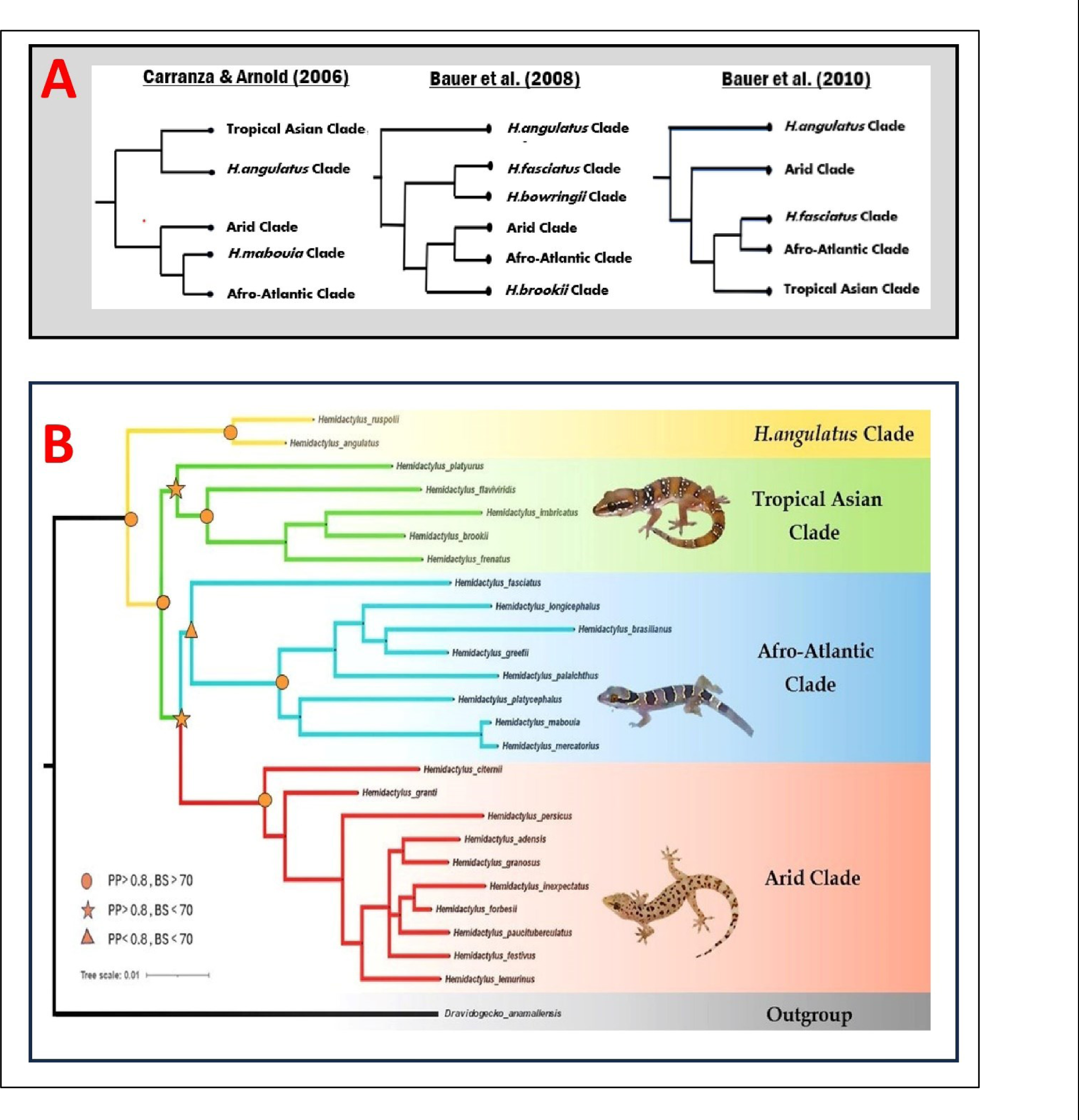
Reduced representation phylogeny of *Hemidactylus* with the resolved clade positions. **Fig 1**- A) The topologies retrieved in the earlier studies, B) The topology retrieved in this study using the nuclear dataset. PP- Posterior Probability, BS- Bootstrap support values.

**Figure 2.**
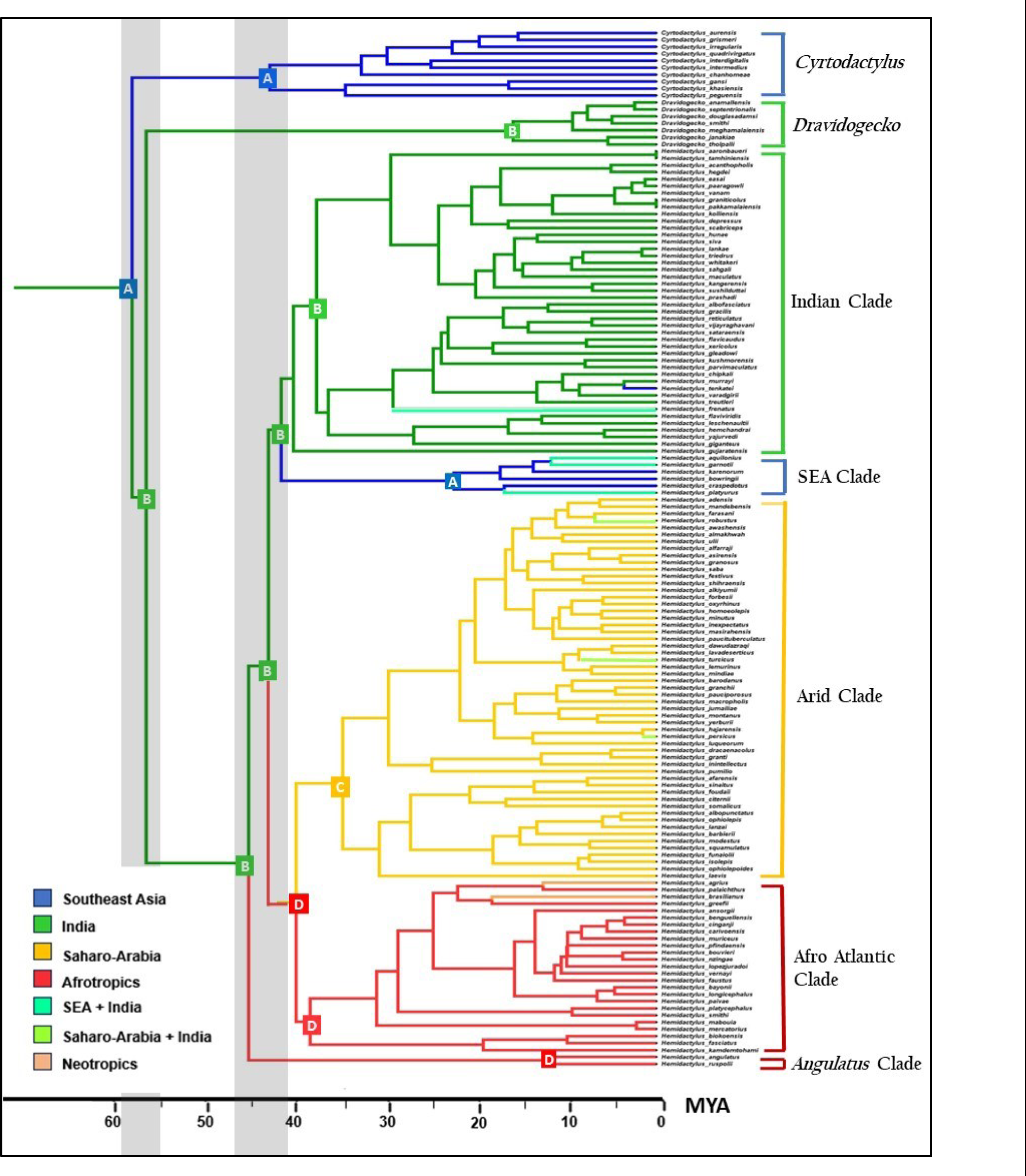
Ancestral area estimation using BioGeoBEARS. **Fig 2**- The result obtained from the BioGeoBEARS analysis for the DEC+j M0 model. The root node of *Hemidactylus* and *Dravidogecko* shows an Indian origin.

Moreover, despite the plethora of studies on *Hemidactylus*, its biogeographic origins remain unclear largely due to conflicting phylogenies and its extensive distribution. Earlier investigations suggested a Southeast Asia origin (Carranza & Arnold, 2006), but these studies overlooked the presence of the endemic Indian sister genus *Dravidogecko*, which raises the possibility of an Indian origin for both genera, and an Out-of-India dispersal scenario for *Hemidactylus*. On the other hand, Africa is the center of diversification for *Hemidactylus* (Smid et al., 2020) and thus an African origin also cannot be overlooked since various studies have linked the centers of diversity to the origin of the groups (Briggs, 2000; Chala et al., 2011; Dupin et al., 2017).

Here, we present the most comprehensive phylogeny of *Hemidactylus* to date with 132 out of the 192 extant species, to resolve uncertainties pertaining to inter-clade relationships. Using a time-calibrated phylogeny, we further reconstruct the historical biogeography of the group to resolve its long-disputed origins and to understand the direction and frequency of dispersal events that may have occurred. Our analysis uses a time-stratified approach in conjunction with plate tectonics information to account for changing distances between landmasses across deep time. We explicitly model and test the Indian origin hypothesis for the genus and compare it with the possibility of either a Saharo-Arabian, an Afrotropical, or a Southeast Asian origin.

## Materials and Methods

### Data acquisition

To determine the higher-level relationships within Hemidactylus, we first downloaded available sequences for six nuclear markers-Recombination Activation Gene 1 (RAG1), Recombination Activation Gene 2 (RAG2), Acetylcholinergic receptor M4 (ACM4), Oocyte Maturation Factor (C-mos), Melanocortin receptor 1 (mc1r), and Phosducin (PDC) from NCBI GenBank (Clark et al., 2016) since nuclear markers are shown to be more informative for deeper nodes (Springer et al., 2001). We generated a reduced-representation dataset consisting of 25 *Hemidactylus* species belonging to the five broad clades defined by Carranza & Arnold (2006) — Tropical Asian clade, Arid clade, Afro-Atlantic clade, *H.angulatus* clade, and *H.mabouia* clade. We included *Dravidogecko anamallensis* as a sister group to our ingroup and *Cyrtodactylus ayeyarwadyensis* as the outgroup. Species were chosen based on the availability of sequences for at least 3 out of the 6 chosen markers (See supplementary table 1 for the details regarding the downloaded sequences). Individual gene alignments were built using the MUSCLE algorithm (Edgar, 2004) in MEGA v.6 (Tamura et al., 2013) using the default settings and were visually inspected for the presence of stop codons using the standard genetic code. The genes were then concatenated to obtain a final dataset of 2951 base pairs (bp). We used PartitionFinder v. 1.1.1 (Lanfear et al., 2012) to partition the dataset by gene and deduce appropriate models of sequence evolution for each partition. The best-fit scheme derived by PartitionFinder was found to be a separate partition per gene with a GTR+I+G model for all the partitions.

### Phylogenetic reconstruction

Phylogenetic reconstruction was carried out using maximum likelihood (ML) as well as Bayesian inference (BI) methods using the obtained partitioning scheme. The ML tree was constructed in RAxML v. 8.0.26 (Stamatakis, 2014), implemented in raxmlGUI v. 1.3.1 (Silvestro & Michalak, 2012) with 1000 thorough bootstrap replicates and GTR+I+G model for all the genes. The Bayesian tree was constructed in MrBayes v.3.2.5 (Ronquist et al., 2012) with the GTR+I+G model. The program was allowed to run for one million generations with trees sampled every 1000^th^ generation, following which the first 25% of the trees were discarded as burn-in. The convergence of the run was ensured by evaluating the standard deviation of split frequencies (<0.01) and ESS values for all the parameters (>200) using the program Tracer V.1.6 (Rambaut et al.,2014).

Additionally, a multispecies coalescent tree was also constructed using *BEAST (Ogilvie et al., 2017) implemented in BEAST v.2.7.6 (Bouckaert et al., 2019) since each of the nuclear genes has an independent evolutionary history. For the coalescent approach, we dropped the mc1r gene as it had very little representation from the Tropical Asian and the Afro-Atlantic clades. The site models and tree models were kept unlinked and the appropriate site model for each gene was estimated using BEAST model test. We unlinked the clock model for each locus under the relaxed lognormal clock with the clock rate set to 0.00076 based on the nuclear substitution rate given for all squamates by Perry et al. (2018). The population function was set at “Linear with constant root” since we did not have an estimate of the effective population size at the root. BEAST was allowed to run 200 million generations with the trees sampled every 10000^th^ generation, following which the ESS values were evaluated using Tracer. A burn-in of 20% was employed to obtain the final Maximum Clade Credibility (MCC) tree. The MCC tree was rooted using *Cyrtodactylus ayeyarwadyensis* following Lajmi & Karanth (2020).

### Time calibration and divergence dating

To obtain a time-calibrated global phylogeny of *Hemidactylus* and *Dravidogecko* for biogeographic reconstruction, we downloaded available sequences for three mitochondrial genes −12S rRNA (12S), Cytochrome-b (Cyt-b), and NADH-dehydrogenase 2 (ND2) from NCBI Genbank (Clark et al., 2016). We also retained PDC and RAG1 since the maximum species had data for these nuclear markers. We concatenated these five markers to obtain a supermatrix with 132 out of 192 *Hemidactylus* species, and 7 out of 8 *Dravidogecko* species (See supplementary table 2 for details regarding downloaded sequences). The dataset was separated by gene and codon position and PartitionFinder was used to estimate the appropriate partitioning scheme and model of sequence evolution to be used as input for BEAST.

We used the program BEAST v.2.6.7 to reconstruct the chronogram. We used the unlinked site model GTR+I+G4 for the mitochondrial genes and TN93 +I +G4 for the nuclear genes according to the output given by PartitionFinder. The clock rates were estimated under the relaxed clock lognormal model. The species were constrained to belong to respective clades based on prior studies (Lajmi et al.,2019; Ceriaco et al.,2020; Smid et al.,2020), whereas the deeper clades were constrained based on the phylogeny obtained in the previous section. Secondary calibrations were used to constrain node ages in the divergence dating analyses based on prior studies using a normal distribution (Chaitanya et al.,2019; Lajmi et al.,2019). The divergence between *Cyrtodactylus* and *Hemidactylus*+*Dravidogecko* was set at a mean age of 60 mya (CI: 50–70 mya), and the split between *Hemidactylus* and *Dravidogecko* at a mean age of 58 mya (CI: 65–50 Mya). The crown age of *Hemidactylus* was given a mean age of 42mya (CI: 38-48mya).

BEAST was allowed to run for 200 million generations with sampling done every 10000^th^ generation under the Yule model of speciation due to the absence of fossil data in our supermatrix. The effective sample size (ESS) values were evaluated to be above 200 using the program TRACER v.1.7. The initial 25% of trees were discarded as burn-in, and the MCC tree was determined using the program TREEANNOTATOR 2.6.4 (Bouckaert et al.,2019).

### Biogeographic reconstruction

The chronogram from the BEAST analysis was used to reconstruct the ancestral areas of *Hemidactylus* and *Dravidogecko* using the R package BioGeoBEARS (Matzke, 2013). The terminal taxa were coded to be present in one or more of the following biogeographic units based on the distribution data obtained from the GBIF database and the updated zoogeographic realms presented by Holt et al. (2013): A= Southeast East Asia, B= Indian subcontinent, C=Saharo-Arabia, D = Afrotropics, and E=Neotropics (Fig 3). Species that were introduced via human mediation in certain regions were excluded from those biogeographic units. The analysis was time-stratified to account for the varying distances between the different land masses over a geological time scale. We used the paleomap dataset (Scotese, 2002) integrated into GPlates plate tectonics visualization software (Müller et al., 2018) to measure the distances between the biogeographic units between 65 mya– 0 mya following the methods of Biswas et al. (2023). The distances measured were then used to create a time-stratified dispersal multiplier matrix by assigning probability values for dispersal between landmasses. Although certain *Hemidactylus* geckos have been shown to undergo transoceanic dispersal, the probability of such long-range dispersals between unconnected landmasses is lesser compared to geodispersal between connected landmasses. Thus, the dispersal multiplier values ranged from 0.99 for landmasses that were adjacent with a land connection, to 0.001 for distant landmasses that were not connected by land. To make the analysis biologically and geologically realistic, we also created a time-stratified areas-adjacency matrix, where closer areas were coded as 1 (adjacent) and farther areas as 0 (not adjacent) to elucidate the combinations of areas allowed where species can simultaneously occur.

**Figure 3.**
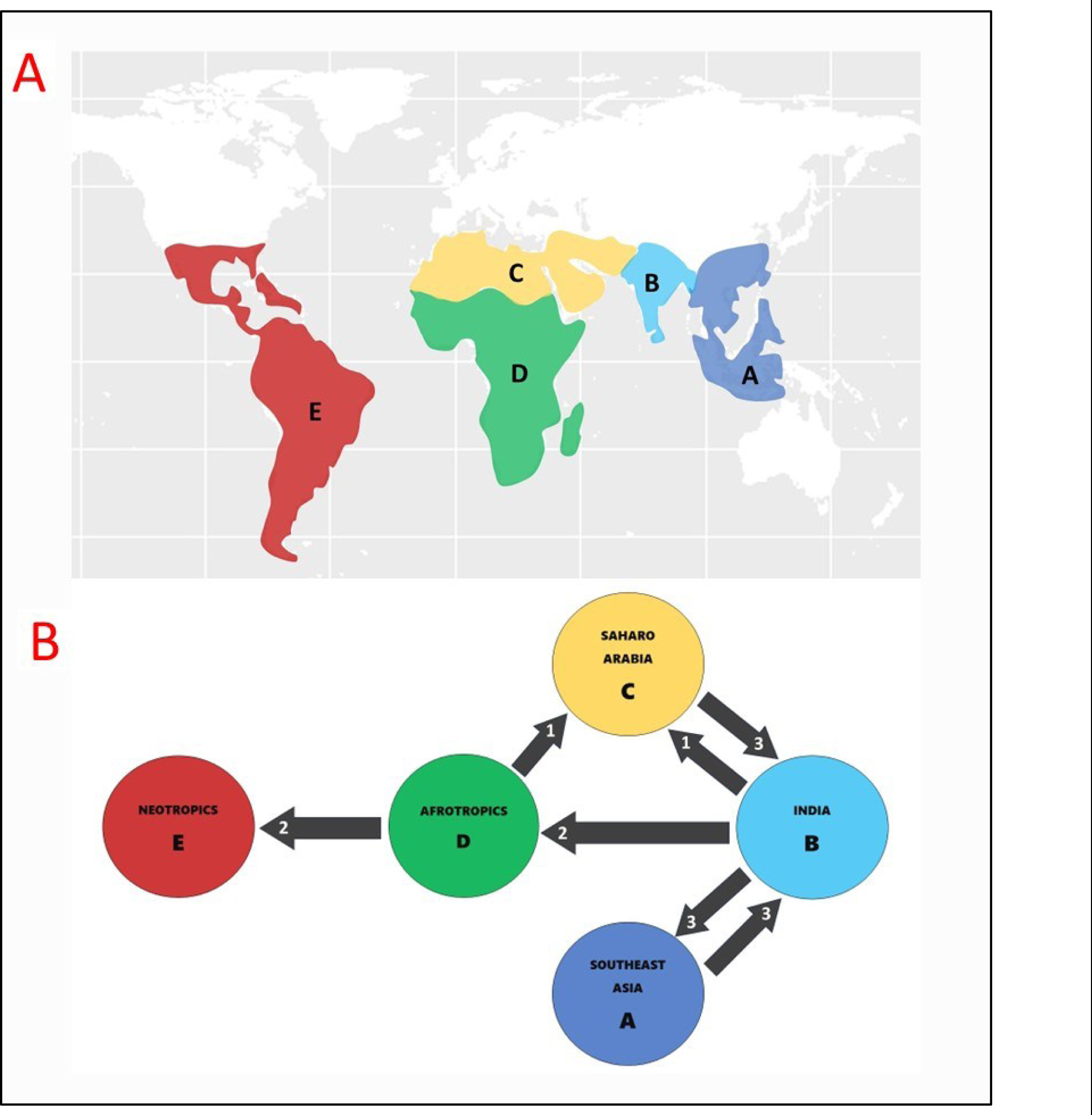
The frequency and directionality of dispersal events within *Hemidactylus*. **Fig 3**- A) The various biogeographical areas coded in this study, B) Dispersals events into and out of the various landmasses. India is shown to be a hub for dispersals with maximum dispersal events.

To rigorously examine the competing hypotheses concerning the origin of *Hemidactylus*, we developed four distinct alternative models by adjusting the dispersal multiplier matrix. The symmetrical dispersal matrix created based on the distance between landmasses was considered to be M0, which does not assume any specific center of origin. The M1 model was designed to reflect an “Indian Origin” for *Hemidactylus*. The dispersal probabilities from India to all the other biogeographic units were kept high (Ranging from 0.75 to 0.95 based on the distances between the drifting continental plates). In contrast, the dispersal probabilities for the “Into-India” scenarios were kept extremely low (0.001) to reflect the Indian origin. The M2 model proposed a Southeast Asian origin, consistent with previous studies (Carranza & Arnold, 2006). Here, dispersal probabilities from South East Asia to other regions were higher (0.75-0.95), while probabilities for dispersal into South East Asia were set to a minimal value (0.001). The M3 model depicted a “Saharo-Arabian” origin, and M4 an “Afrotropical” origin, where dispersal probabilities from Middle and East Africa to other regions were high (0.75-0.95). In contrast, probabilities for dispersal from India, South East Asia, and the Neotropics into Africa and the Middle East were kept low (0.001).

To comprehensively evaluate these alternative origin scenarios, ancestral area reconstructions were performed for each of the alternative models under two basic BioGeoBEARS models — “DEC” (Dispersal-Extinction-Cladogenesis model, Ree & Smith, 2008), and “DIVALIKE” (A likelihood version of Dispersal–Vicariance Analysis model, Roquist, 1997). We did not utilize the “BAYAREALIKE” model since it assumes a static geological history (Landis et al., 2013). These two models were also run with an additional “j” parameter that incorporates founder event speciation or long-range dispersals (Matzke, 2013). We selected the model that was the best fit for the data based on corrected Akaike information criterion (AIC) scores (see Table 1).

**Table 1.** Model fitting using BioGeoBEARS. Table 1-The various models created for alternative hypothesis testing using BioGeoBEARS. The models showing the lowest AIC scores have been highlighted in red.

| Sr. No | BioGeoBEARS iteration | Model | DEC | DEC+J | DIVALIKE | DIVALIKE+J |
| --- | --- | --- | --- | --- | --- | --- |
| 1 | With sister groups | **M0** | 258.88 | **164.4** | 273.54 | 166.66 |
| 2 |  | M1 (Indian origin) | 331.56 | 218 | 358.8 | 237.58 |
| 3 |  | M2 (Southeast Asian origin) | 392.18 | 221.1 | 423.03 | 223.22 |
| 4 |  | M3 (Saharo-Arabian origin) | 398.28 | 256.8 | 423.64 | 268.2 |
| 5 |  | M4 (Afrotropical origin) | 426.82 | 245.3 | 461.08 | 256.74 |
| 1 | Without sister groups | **M0** | 150.18 | **142.9** | 145.66 | 144.26 |
| 2 |  | M1 (Indian origin) | 222.16 | 201.6 | 227.62 | 220.66 |
| 3 |  | M2 (Southeast Asian origin) | 197.74 | 169.8 | 198 | 178.24 |
| 4 |  | M3 (Saharo-Arabian origin) | 234.1 | 233.3 | 243.14 | 244.64 |
| 5 |  | M4 (Afrotropical origin) | 249.42 | 228.6 | 266 | 239.88 |

In our BioGeoBEARS analysis, we coded Saharo-Arabia and Afrotropics as two distinct areas based on the classifications of Holt et al (2013), since these areas show variations in climate and vegetation (henceforth termed as the “five area” reconstruction). But Africa and the Middle East were geologically a single landmass till the African plate rifted apart from the Arabian plate during the mid-Oligocene to early Miocene approximately 31-23 mya (Smid et al., 2013). We thus also performed an additional BioGeoBEARS run after coding these two landmasses to be a single biogeographic unit (henceforth termed as the “four area” reconstruction).

Since ancestral area reconstructions are sensitive to the choice of the outgroups (Stevens & Heesy, 2006), especially when the ingroup is cosmopolitan and the sister group is range-restricted, we analyzed all the above-mentioned models by first including *Dravidogecko* and *Cyrtodactylus* in the phylogeny followed by their exclusion, and subsequently compared all these reconstructions.

## Results

### Phylogeny reconstruction & divergence dating

The reduced representation dataset included 25 ingroup and two outgroup species. The total concatenated sequence length for RAG1 (848 bp), RAG2(366 bp), ACM4(386 bp), C-mos (358 bp), PDC (327 bp), and mc1r (666 bp) consisted of 2951 sites of which 256 were parsimony informative.

The topologies were similar in all three tree-building methods barring minor incongruencies in relationships within the clades, but these were not relevant since our focus was on the resolution of the higher-level relationships. However, node supports were consistently lower for the coalescent tree (See supplementary material 2). Due to the lack of available nuclear data, we only had one representative from the Southeast Asian species (*H.platyurus*) which was sister to a clade comprising species from India with high support (PP=0.98), thus ascertaining the monophyly of the Tropical Asian clade. Our phylogeny also recovered branches leading to the Arid and *H. angulatus* clades with very high support (PP > 0.95). The *H.angulatus* clade was retrieved as the sister group to the rest of *Hemidactylus* with strong support (PP=1). While *H.mabouia* was nested within the Afro-Atlantic Clade, *H.fasciatus* was recovered as a sister to the Afro-Atlantic clade, but with moderate support(PP=0.72, see Fig1).

The chronogram suggested that the first split within *Hemidactylus* occurred between the ancestor of the *H. anuglatus* clade and the rest of *Hemidactylus* approximately 45 mya. The second split occurred between the Arid clade + Afro-Atlantic/*H.mabouia*/*H.fasciatus* clade and the Tropical Asian clade around 42 mya, and the Arid Clade further split from the Afro-Atlantic clade approximately 38 mya (See Fig 2).

### Biogeographic reconstructions

Out of the five models created to test the biogeographic origins of the genus, the DEC+j M0 model which included the distance matrix, an areas adjacency matrix, and a symmetrical dispersal probability multiplier received the lowest AIC score. This was followed by the DIVALIKE+j M0 model (Table 1).

The biogeographic reconstruction assigned Southeast Asia to the ancestral node of *Hemidactylus, Dravidogecko,* and *Cyrtodactylus* at approximately 58 mya (Fig 2). Importantly, the reconstruction at the ancestral node of *Hemidactylus* and *Dravidogecko* suggested an Indian origin for both these genera. The crown group of *Hemidactylus* was also estimated to be in the Indian subcontinent, lending support to the Indian origin. Within *Hemidactylus*, there were three major dispersals out of India corresponding to the formation of the major global clades. The first dispersal from India to Afro-Arabia happened around 45 mya which led to the formation of the *H.angulatus* clade while the second dispersal around 42 mya gave rise to the Arid clade and the Afro-Atlantic clade. The third major dispersal event transpired from India to Southeast Asia around 22 mya post the suturing of the Indian plate with Asia, leading to the Southeast Asian/*bowringii* subclade within the Tropical Asian clade. Within the Afro-Atlantic clade, there were multiple independent colonisations of South America in the last 20 million years The reconstruction also indicated back dispersals and range extensions into the Indian subcontinent within a few species ——*H.aquilonius*, *H.garnotii*, and *H.platyurus* expanded into Northeast India from Southeast Asia, and *H.robustus, H.turcicus*, and *H.persicus* dispersed into western Indian from the Saharo-Arabian region (Fig 2 and Fig 3)..

Dec M0 model emerged as the best model for the biogeographic reconstructions after coding Africa and the Middle East to be a single biogeographic unit (the “four area” reconstruction). This model also showcased India at the node of *Hemidactylus* and *Dravidogecko*, but the results demonstrated a widespread common ancestor in Southeast Asia and India at the node of *Hemidactylus* + *Dravidogecko* and *Cyrtodactylus*. The crown group of *Hemidactylus* was reported to be widespread in India and Africa. These reconstructions are unrealistic since India, Southeast Asia, and Africa were well separated from each other during this period. Therefore we report only the “five area” reconstruction in the paper (refer to supplementary material 4 for details regarding the “four area” BioGeoBEARS result).

The biogeographic analysis excluding *Cyrtodactylus* and *Dravidogecko* from the analysis gave a starkly contrasting reconstruction at the ancestral node of *Hemidactylus*. The best model was the DEC+j M0 followed by the DIVALIKE+j M0. Both these models suggested an Afrotropical origin for the genus, with dispersals to India and Saharo-Arabia around 43 mya and 38 mya respectively (See supplementary material 5).

## Discussion

### Resolution of inter-clade relationships

A pioneering study by Carranza & Arnold (2006) endeavoured to resolve the global phylogeny of *Hemidactylus* using two mitochondrial markers. They retrieved five main clades corresponding to major world regions. In their phylogeny, the *H.angulatus* clade was sister to the Tropical Asian clade; the Afro-Atlantic clade was a sister to the Arid clade; whereas *H. fasciatus* could not be allotted to any clade. Notably, they found support for the monophyly of the Tropical Asian group with two reciprocally monophyletic subgroups; one with Indian species and the other with Southeast Asian species. However, Bauer et al. (2008) proposed that Tropical Asian *Hemidactylus* were not monophyletic. The inter-clade relationships also differed between the analyses, and the position of *H.fasciatus* remained unresolved. The phylogeny of Bansal and Karanth (2010) included additional species from the Indian radiation and retrieved the Tropical Asian clade (sensu Carranza & Arnold, 2006) using the mitochondrial dataset, but not in the nuclear phylogeny. Finally, Bauer et al. (2010) retrieved the Tropical Asian clade using two nuclear and mitochondrial markers each, although here it was sister to the Afro-Atlantic/ *H.mabouia* clade. The *H. angulatus* clade was sister to the rest of the *Hemidactylus* species, and the placement of *H.fasciatus* was still not well supported (See Fig 1) These inconsistencies in the topologies served as a limitation for the studies attempting to investigate the macroevolutionary dynamics of this group.

Our phylogeny built using the nuclear dataset resolved these conflicting inter-clade relationships within *Hemidactylus* with good support and integrated the results from the earlier studies. The Indian and the Southeast Asian species together constituted the larger Tropical Asian clade. The results of our reconstruction thus confirmed the monophyly of the Asian *Hemidactylus*, supporting the findings of Carranza & Arnold (2006) and Bauer et al. (2010).

*H. fasciatus* and *H. mabouia* are both believed to have evolved in Sub-Saharan Africa (Ceriaco et al.,2020; Agarwal et al.,2021). This scenario was supported in the phylogeny where *H. fasciatus* and *H. mabouia* fell within the Afro-Atlantic clade that encompassed the Sub-Saharan region in our analysis. The neotropical species were also nested within the Afro-Atlantic clade, alluding to multiple independent colonisations of the neotropics by the African lineages. Further, the placement of the *H.angulatus* clade as the sister for all other *Hemidactylus*, consistent with Bauer et al. (2010), was retrieved with high support by all the tree-building methods. Africa is home to two *Hemidactylus* clades, with the Arid clade situated in the Saharo-Arabian region, and the Afro-Atlantic clade in Central and Southern Africa. The existence and preservation of these distinct clades in Africa, despite the absence of prominent geographic barriers, could be attributed to the “eco-climatic” transitions between the drier regions in northern Africa and the more humid, equatorial areas in central Africa (Happold and Lock, 2013) which may have led to the split between the lineages leading to the two clades.

### Importance of including appropriate sister groups in biogeographic reconstructions

Earlier studies postulated that *Hemidactylus* originated in Southeast Asia and after the fusion of the Indian plate with the Eurasian plate, some lineages dispersed into India and Africa where they subsequently underwent radiations to form today’s geographically delineated clades (Carranza & Arnold, 2006; Bansal & Karanth 2013). However, none of these studies employed explicit biogeographic analyses to test these assumptions. Our study contradicted these results and demonstrated an Indian origin for *Hemidactylus* using extensive global sampling, alternative models for hypothesis testing, incorporation of plate tectonic information, and time stratification. Before the Indian endemic *Dravidogecko* was assigned as the sister genus to *Hemidactylus*, the closest relative of *Hemidactylus* based on morphology was proposed to be the largely Asian *Cyrtodactylus* (Carranza & Arnold, 2006). Thus, including the more distantly related group might have shown a Southeast Asian origin for the genus. Similarly, excluding *Dravidogecko* from the analyses yielded an African origin. This was probably because the *H.angulatus* clade showing the earliest divergence from the rest of the species is distributed in Africa, and most of the species are concentrated within this region. We thus suggest the importance of including the sister group (especially if it is range-restricted) following the latest taxonomic revisions to prevent inaccurate estimations of ancestral areas in the case of widely distributed groups.

### Biogeographic reconstruction and Indian origin

The biogeographic reconstruction placed the ancestor of *Hemidactylus*, *Dravidogecko*, and *Cyrtodactylus* in Southeast Asia, following which the lineage leading to *Hemidactylus* and *Dravidogecko* dispersed into India between 58-56 mya. The nature of this dispersal is ambiguous since hypothesized land bridges between the drifting Indian plate and Southeast Asia could have enabled geodispersal (Ali & Aitchison, 2008; Grismer et al.,2016; Yuan et al., 2019). However, Klaus et al. (2016) stated that the dispersal events between India and Asia before 44 mya were likely to be transmarine, restricted to taxa that could surmount marine barriers. This assumption, coupled with the propensity of geckos to undergo transoceanic dispersal through saltwater tolerance, indicated that an ancestral population likely “jumped” onto the drifting Indian plate from Southeast Asia and subsequently diverged into *Hemidactylus* and *Dravidogecko* in India.

The unique geological origin of the Indian subcontinent conferred upon it a multifaceted role in the community assemblage of the Indian and Southeast Asian biota. Most of the current Indian tetrapod lineages were the consequences of the dispersals and colonisations from Southeast Asia to India (Karanth 2021). The Out-of-India dispersals shown till now could be classified broadly into two categories— (A) India acted as a “biotic ferry” to transport ancient Gondwanan forms such as caecilians (Zhang & Wake, 2009), blindsnakes (Sidharthan & Karanth,2021) and centipedes (Joshi & Karanth, 2011) into Southeast Asia, and (B) Various groups that originated in one region utilised the drifting Indian plate as a “stepping stone” to disperse to another region, such as tarantulas (Biswas et al.,2023), Natatanuran frogs (Yuan et al.,2019), and Dipterocarps (Bansal et al.,2022). However, examples of non-Gondwanan groups originating in the insular Indian plate and further dispersing outwards were uncommon, some exceptions being Ranid frogs (Bossuyt & Milkinovitch, 2001), rice fishes (Yamahira et al.,2021), and Heterometrinae scorpions (Loria & Pendini, 2020). Notably, no such examples have been reported for Indian squamates.

The origin of a widespread and focal genus such as *Hemidactylus* in India presented evidence of one such rare instance of non-Gondwanan radiation that originated in the Indian landmass and further underwent “Out-Of-India” dispersal. The intricate biogeography of *Hemidactylus*, involving dispersals into Africa and Asia, as well as back dispersals into India, signified a novel and unique biogeographic pattern in the context of Indian squamates. Krause & Maas (1990) suggested that “among early Tertiary large landmasses, the Indian subcontinent is unique in its combination of having been in the right places at the right times to provide for the development and the subsequent disembarking of several new higher taxa of mammals”. Our results expanded upon this pattern and indicated that the emergence of other vertebrate lineages may also have origins in India, notwithstanding significant isolation and widespread volcanism.

## Supporting information

Coalescent phylogeny

BioGeoBEARS 5 area result

BioGeoBEARS 4 area result

BioGeoBEARS result without sister groups

## Acknowledgments

The authors would like to sincerely thank R. Chaitanya for his insightful comments regarding the manuscript. We would also like to acknowledge the Institute of Eminence (IOE) scheme for funding our research.

## Notes

### Competing Interest Statement

The authors have declared no competing interest.

