## Supplemental tables 1 and 2 for "Globetrotting geckos: Historical biogeography suggests an Indian origin and “Out-Of-India” dispersal for the cosmopolitan *Hemidactylus* geckos"

**Supplementary Table 1**

Details regarding the molecular dataset utilized for building the reduced representation phylogeny

| No. | Species | Clade | RAG1 | PDC | RAG2 | ACM4 | CMOS | mc1r |
|---|---|---|---|---|---|---|---|---|
| 1 | *Hemidactylus brasilianus* | Afro-Atlantic clade | EU268290.1 | EU268320.1 | HQ426439.1 | HQ426346.1 | HQ426523.1 | **NA** |
| 2 | *Hemidactylus greefii* | Afro-Atlantic clade | MT613940.1 | EU268338.1 | HQ426459.1 | HQ426367.1 | HQ426542.1 | **NA** |
| 3 | *Hemidactylus longicephalus* | Afro-Atlantic clade | MZ617049.1 | MN843914.1 | HQ426460.1 | HQ426369.1 | HQ426544. | **NA** |
| 4 | *Hemidactylus mercatorius* | Afro-Atlantic clade | MN853057.1 | MN843920.1 | **NA** | **NA** | AY863046.1 | **NA** |
| 5 | *Hemidactylus palaichthus* | Afro-Atlantic clade | EU268307.1 | EU268337.1 | HQ426464.1 | HQ426373.1 | HQ426548.1 | **NA** |
| 6 | *Hemidactylus platycephalus* | Afro-Atlantic clade | JQ073246.1 | **NA** | KC819045.1 | **NA** | KC818776.1 | KC818926.1 |
| 7 | *Hemidactylus ruspolii* | Afro-Atlantic clade | **NA** | **NA** | KC819052.1 | **NA** | KC818782.1 | KC818931.1 |
| 8 | *Hemidactylus angulatus* | Angulatus clade | HM559686.1 | EU268336.1 | **NA** | **NA** | KC818747.1 | KC818903.1 |
| 9 | *Hemidactylus citernii* | Arid Clade | MN538124.1 | KU749871.1 | MN538167.1 | KU749854.1 | MN538044.1 | MN538089.1 |
| 10 | *Hemidactylus adensis* | Arid Clade | KP238242.1 | OL653910.1 | KP238233.1 | **NA** | KP238253.1 | KP238245.1 |
| 11 | *Hemidactylus festivus* | Arid Clade | KC818961.1 | KU749874.1 | KC819024.1 | KU749839.1 | JQ957125.1 | KC818909.1 |
| 12 | *Hemidactylus forbesii* | Arid Clade | KU955730.1 | KU749876.1 | KU749933.1 | KU749845.1 | KU749861.1 | JQ982645.1 |
| 13 | *Hemidactylus granosus* | Arid Clade | KF647595.1 | OL653954.1 | OL653863.1 | KU749847.1 | OL653789.1 | OL653897.1 |
| 14 | *Hemidactylus inexpectatus* | Arid Clade | KU955733.1 | KU749880.1 | JQ957426.1 | KU749843.1 | JQ957140.1 | JQ957273.1 |
| 15 | *Hemidactylus lemurinus* | Arid Clade | KC818978.1 | KU749882.1 | JQ957423.1 | KU749842.1 | JQ957137.1 | KC818920.1 |
| 16 | *Hemidactylus persicus* | Arid Clade | KU749926.1 | EU268346.1 | KU749938.1 | KU749849.1 | KC818775.1 | KC818924.1 |
| 17 | *Hemidactylus paucituberculatus* | Arid Clade | KU955738.1 | KU749886.1 | JQ957433.1 | KU749840.1 | JQ957150.1 | JQ957284.1 |
| 18 | *Hemidactylus granti* | Arid Clade | KU955732.1 | KU749877.1 | KC819032.1 | KU749851.1 | KC818759.1 | JQ982650.1 |
| 19 | *Hemidactylus fasciatus* | *Fasciatus* clade | JQ945311.1 | JQ945379.1 | JQ945470.1 | JQ945683.1 | JQ945577.1 | **NA** |
| 20 | *Hemidactylus mabouia* | *Mabouia* Clade | MW790458.1 | MW733817.1 | HQ426461.1 | HQ426370.1 | KC818771.1 | KC818922.1 |
| 21 | *Hemidactylus platyurus* | Tropical Asian | HM559685.1 | MW133495.1 | KY366182.1 | HQ426353.1 | HQ426530.1 | **NA** |
| 22 | *Hemidactylus brookii* | Tropical Asian | EU268305 | EU268335 | KJ131425.1 | **NA** | KC735118.1 | **NA** |
| 23 | *Hemidactylus flaviviridis* | Tropical Asian | MN482264.1 | EU268324.1 | KY366189.1 | HQ426366.1 | HQ426541.1 | KC818911.1 |
| 24 | *Hemidactylus imbricatus* | Tropical Asian | EU268293 | EU268323 | HQ426506.1 | HQ426416.1 | HQ426587.1 | **NA** |
| 25 | *Hemidactylus frenatus* | Tropical Asian | HM559695.1 | EU268328.1 | EF534982.1 | EF534897.1 | EF534940.1 | **NA** |
| 26 | *Dravidogecko anamallensis* | | KC735090.1 | MN520287.1 | **NA** | **NA** | KC735117.1 | **NA** |
| 27 | *Cyrtodactylus ayeyarwadyensis* | | EU268287.1 | EU268317.1 | JQ945443.1 | JQ945656.1 | JQ945550.1 | NA |

**Supplementary Table 2**

Details regarding the molecular dataset utilized for building the BEAST chronogram

| No | Species | Clade | 12S | CytB | ND2 | RAG1 | PDC |
|---|---|---|---|---|---|---|---|
| 1 | *Hemidactylus agrius* | Afro-Atlantic clade | DQ120433.1 | DQ120262.1 | **NA** | **NA** | **NA** |
| 2 | *Hemidactylus ansorgii* | Afro-Atlantic clade | **NA** | **NA** | MN843756.1 | MN853036.1 | MN843904.1 |
| 3 | *Hemidactylus bayonii* | Afro-Atlantic clade | **NA** | **NA** | MZ616911.1 | MZ617017.1 | MN843905.1 |
| 4 | *Hemidactylus benguellensis* | Afro-Atlantic clade | **NA** | **NA** | MZ616926.1 | MZ617031.1 | MN843910.1 |
| 5 | *Hemidactylus biokoensis* | Afro-Atlantic clade | **NA** | EU268371 | **NA** | **NA** | **NA** |
| 6 | *Hemidactylus boavistensis (Subsp. Bouvieri)* | Afro-Atlantic clade | **NA** | EU730675.1 | **NA** | **NA** | **NA** |
| 7 | *Hemidactylus bouvieri* | Afro-Atlantic clade | AF324812.1 | DQ120252.1 | **NA** | **NA** | **NA** |
| 8 | *Hemidactylus brasilianus* | Afro-Atlantic clade | DQ120428.1 | EU268383.1 | EU268351.1 | EU268290.1 | EU268320.1 |
| 9 | *Hemidactylus carivoensis* | Afro-Atlantic clade | **NA** | **NA** | MZ616927.1 | MZ617032.1 | **NA** |
| 10 | *Hemidactylus cinganji* | Afro-Atlantic clade | **NA** | **NA** | MZ616931.1 | MZ617039.1 | **NA** |
| 11 | *Hemidactylus faustus* | Afro-Atlantic clade | **NA** | **NA** | MZ616933.1 | MZ617041.1 | **NA** |
| 12 | *Hemidactylus greefii* | Afro-Atlantic clade | DQ120413.1 | EU268401.1 | MT613970.1 | MT613940.1 | EU268338.1 |
| 13 | *Hemidactylus kamdemtohami* | Afro-Atlantic clade | **NA** | **NA** | MN843790.1 | MN853048.1 | MN843911.1 |
| 14 | *Hemidactylus longicephalus* | Afro-Atlantic clade | DQ120416.1 | DQ120246.1 | MZ616948.1 | MZ617049.1 | MN843914.1 |
| 15 | *Hemidactylus lopezjuradoi* | Afro-Atlantic clade | EU730639.1 | EU730650.1 | **NA** | **NA** | **NA** |
| 16 | *Hemidactylus mercatorius* | Afro-Atlantic clade | **NA** | **NA** | MN843845.1 | MN853057.1 | MN843920.1 |
| 17 | *Hemidactylus muriceus* | Afro-Atlantic clade | **NA** | **NA** | MN843847.1 | MN853059.1 | MN843922.1 |
| 18 | *Hemidactylus nzingae* | Afro-Atlantic clade | **NA** | **NA** | MZ616975.1 | MZ617095.1 | |
| 19 | *Hemidactylus paivae* | Afro-Atlantic clade | **NA** | **NA** | MZ616984.1 | MZ617104.1 | MN843926.1 |
| 20 | *Hemidactylus palaichthus* | Afro-Atlantic clade | DQ120434.1 | EU268400.1 | EU268368.1 | EU268307.1 | EU268337.1 |
| 21 | *Hemidactylus pfindaensis* | Afro-Atlantic clade | **NA** | **NA** | MZ616990.1 | MZ617110.1 | **NA** |
| 22 | *Hemidactylus platycephalus* | Afro-Atlantic clade | KC818692.1 | KC818845.1 | MW790401.1 | JQ073246.1 | **NA** |
| 23 | *Hemidactylus principensis* | Afro-Atlantic clade | **NA** | **NA** | MT613971.1 | MT613941.1 | **NA** |
| 24 | *Hemidactylus ruspolii* | Afro-Atlantic clade | KC818706.1 | KC818860.1 | **NA** | **NA** | **NA** |
| 25 | *Hemidactylus smithi* | Afro-Atlantic clade | KC818715.1 | KC818870.1 | **NA** | KC818996.1 | **NA** |
| 26 | *Hemidactylus vernayi* | Afro-Atlantic clade | **NA** | **NA** | MZ616994.1 | MZ617113.1 | **NA** |

| | | | | | | | |
|---|---|---|---|---|---|---|---|
| 27 | *Hemidactylus gramineus* | Afro-Atlantic clade | **NA** | **NA** | MN843869.1 | **NA** | **NA** |
| 28 | *Hemidactylus angulatus* | Angulatus clade | KC818667.1 | DQ120238.1 | HM559620.1 | HM559686.1 | EU268336.1 |
| 29 | *Hemidactylus afarensis* | Arid Clade | **NA** | MN537997.1 | **NA** | **NA** | **NA** |
| 30 | *Hemidactylus albopunctatus* | Arid Clade | KC818657.1 | KC818794.1 | **NA** | KC818952.1 | **NA** |
| 31 | *Hemidactylus barbierii* | Arid Clade | KU955711.1 | MN537998.1 | **NA** | MN538123.1 | **NA** |
| 32 | *Hemidactylus citernii* | Arid Clade | KC818670.1 | MN537999.1 | **NA** | MN538124.1 | KU749871.1 |
| 33 | *Hemidactylus foudaii* | Arid Clade | KC818677.1 | KU955720.1 | **NA** | KC818968.1 | **NA** |
| 34 | *Hemidactylus funaiolii* | Arid Clade | KC818678.1 | KC818827.1 | **NA** | KC818969.1 | **NA** |
| 35 | *Hemidactylus isolepis* | Arid Clade | KC818680.1 | KC818835.1 | **NA** | MN538126.1 | **NA** |
| 36 | *Hemidactylus klauberi* | Arid Clade | **NA** | **NA** | **NA** | **NA** | **NA** |
| 37 | *Hemidactylus laevis* | Arid Clade | **NA** | MN538003.1 | **NA** | MN538138.1 | **NA** |
| 38 | *Hemidactylus lanzai* | Arid clade | **NA** | MN538006.1 | **NA** | MN538140.1 | **NA** |
| 39 | *Hemidactylus modestus* | Arid Clade | DQ120386.1 | DQ120215.1 | **NA** | **NA** | **NA** |
| 40 | *Hemidactylus ophiolepis* | Arid Clade | KC818689.1 | KC818841.1 | **NA** | KC818982.1 | **NA** |
| 41 | *Hemidactylus ophiolepoides* | Arid Clade | **NA** | MN538015.1 | **NA** | MN538145.1 | **NA** |
| 42 | *Hemidactylus sinaitus* | Arid Clade | KC818712.1 | JQ957231.1 | **NA** | MN538128.1 | **NA** |
| 43 | *Hemidactylus somalicus* | Arid Clade | **NA** | MN538018.1 | **NA** | MN538130.1 | **NA** |
| 44 | *Hemidactylus squamulatus* | Arid Clade | KC818737.1 | MN538027.1 | **NA** | KC819005.1 | **NA** |
| 45 | *Hemidactylus achaemenidicus* | Arid Clade | **NA** | **NA** | **NA** | MN538121.1 | **NA** |
| 46 | *Hemidactylus adensis* | Arid Clade | KP238276.1 | KP238268.1 | **NA** | KP238242.1 | OL653910.1 |
| 47 | *Hemidactylus alfarraji* | Arid Clade | KX263484.1 | KX263649.1 | **NA** | KX263546.1 | OL653919.1 |
| 48 | *Hemidactylus alkiyumii* | Arid Clade | JQ957095.1 | JQ957178.1 | **NA** | KC818955.1 | KU749870.1 |
| 49 | *Hemidactylus almakhwah* | Arid Clade | NC_066453.1 | NC_066453.1 | NC_066453.1 | **NA** | **NA** |
| 50 | *Hemidactylus asirensis* | Arid Clade | KX263492.1 | KX263657.1 | **NA** | KX263554.1 | OL653934.1 |
| 51 | *Hemidactylus awashensis* | Arid Clade | KP238269.1 | KP238266.1 | **NA** | KP238240.1 | OL653936.1 |
| 52 | *Hemidactylus barodanus* | Arid Clade | KX263493.1 | KC818814.1 | **NA** | KX263558.1 | **NA** |
| 53 | *Hemidactylus dawudazraqi* | Arid Clade | KC818671.1 | JQ957230.1 | **NA** | KC818959.1 | KU749872.1 |
| 54 | *Hemidactylus farasani* | Arid Clade | NC_066454.1 | OL653737.1 | NC_066454.1 | OL653814.1 | **NA** |
| 55 | *Hemidactylus festivus* | Arid Clade | JQ957097.1 | JQ957179.1 | **NA** | KC818961.1 | KU749874.1 |
| 56 | *Hemidactylus forbesii* | Arid Clade | JQ982785.1 | JQ982893.1 | **NA** | KU955730.1 | KU749876.1 |
| 57 | *Hemidactylus granchii* | Arid Clade | KC818679.1 | KC818828.1 | **NA** | KC818970.1 | **NA** |
| 58 | *Hemidactylus granosus* | Arid Clade | KU955712.1 | KU955718.1 | **NA** | KF647595.1 | OL653954.1 |
| 59 | *Hemidactylus hajarensis* | Arid Clade | JQ957050.1 | JQ957185.1 | **NA** | KC818972.1 | KU749878.1 |
| 60 | *Hemidactylus homoeolepis* | Arid Clade | KJ189812.1 | KJ189775.1 | **NA** | KJ189931.1 | KU749879.1 |
| 61 | *Hemidactylus inexpectatus* | Arid Clade | JQ957065.1 | JQ957205.1 | **NA** | KU955733.1 | KU749880.1 |

| | | | | | | | |
|---|---|---|---|---|---|---|---|
| 62 | *Hemidactylus jumailiae* | Arid Clade | KX263494.1 | KX263658.1 | **NA** | KX263559.1 | **NA** |
| 63 | *Hemidactylus lavadeserticus* | Arid Clade | KC818683.1 | DQ120165.1 | **NA** | KC818976.1 | **NA** |
| 64 | *Hemidactylus lemurinus* | Arid Clade | JQ957061.1 | JQ957199.1 | **NA** | KC818978.1 | KU749882.1 |
| 65 | *Hemidactylus luqueorum* | Arid Clade | JQ957069.1 | JQ957211.1 | **NA** | **NA** | **NA** |
| 66 | *Hemidactylus macropholis* | Arid Clade | KX263500.1 | KX263665.1 | JX041369.1 | KX263564.1 | HQ426203.1 |
| 67 | *Hemidactylus mandebensis* | Arid Clade | NC_066455.1 | NC_066455.1 | NC_066455.1 | KP238236.1 | OL653959.1 |
| 68 | *Hemidactylus masirahensis* | Arid Clade | JQ957063.1 | JQ957202.1 | **NA** | KU955736.1 | KU749884.1 |
| 69 | *Hemidactylus mindiae* | Arid Clade | KC818686.1 | HQ833747.1 | **NA** | KC818989.1 | **NA** |
| 70 | *Hemidactylus minutus* | Arid Clade | KJ189804.1 | KJ189773.1 | **NA** | KJ189922.1 | **NA** |
| 71 | *Hemidactylus montanus* | Arid Clade | KX263476.1 | KX263667.1 | **NA** | KX263580.1 | **NA** |
| 72 | *Hemidactylus oxyrhinus* | Arid Clade | JQ982807.1 | DQ120173.1 | **NA** | KU955737.1 | KU749885.1 |
| 73 | *Hemidactylus pauciporosus* | Arid Clade | DQ120379.1 | DQ120208.1 | **NA** | **NA** | **NA** |
| 74 | *Hemidactylus persicus* | Arid Clade | MG744556.1 | EU268409.1 | EU268377 | KU749926.1 | EU268346.1 |
| 75 | *Hemidactylus pseudoromeshkanicus* | Arid Clade | **NA** | **NA** | **NA** | **NA** | **NA** |
| 76 | *Hemidactylus robustus* | Arid Clade | KX263520.1 | OL653758.1 | HM559644.1 | KP238238.1 | OL653963.1 |
| 77 | *Hemidactylus romeshkanicus* | Arid Clade | MG744524.1 | **NA** | **NA** | **NA** | **NA** |
| 78 | *Hemidactylus saba* | Arid Clade | KF647567.1 | KF647579.1 | **NA** | KF647594.1 | OL653964.1 |
| 79 | *Hemidactylus shihraensis* | Arid Clade | KC818707.1 | KC818862.1 | **NA** | **NA** | **NA** |
| 80 | *Hemidactylus turcicus* | Arid Clade | HQ675924.1 | HQ676054.1 | EU268360.1 | HQ426293.1 | EU268329.1 |
| 81 | *Hemidactylus ulii* | Arid Clade | KF647572.1 | NC_066456.1 | NC_066456.1 | KF647603.1 | OL653968.1 |
| 82 | *Hemidactylus yerburii* | Arid Clade | JQ957085.1 | JQ957235.1 | **NA** | KC819011.1 | **NA** |
| 83 | *Hemidactylus paucituberculatus* | Arid Clade | JQ957073.1 | JQ957217.1 | **NA** | KU955738.1 | KU749886.1 |
| 84 | *Hemidactylus dracaenacolus* | Arid Clade | JQ982781.1 | JQ982889.1 | **NA** | KU955729.1 | KU749873.1 |
| 85 | *Hemidactylus granti* | Arid Clade | JQ982789.1 | JQ982898.1 | **NA** | KU955732.1 | KU749877.1 |
| 86 | *Hemidactylus inintellectus* | Arid Clade | JQ982805.1 | JQ982921.1 | **NA** | KU955734.1 | KU749881.1 |
| 87 | *Hemidactylus pumilio* | Arid Clade | JQ982817.1 | DQ120211.1 | **NA** | KU955739.1 | |
| 88 | *Hemidactylus coalescens* | Fasciatus clade | HM180302.1 | **NA** | **NA** | **NA** | **NA** |
| 89 | *Hemidactylus eniangii* | Fasciatus clade | HM180287.1 | **NA** | **NA** | **NA** | **NA** |
| 90 | *Hemidactylus fasciatus* | Fasciatus clade | HM180291.1 | **NA** | EU268370.1 | JQ945311.1 | JQ945379.1 |
| 91 | *Hemidactylus kyaboboensis* | Fasciatus clade | HM180289.1 | **NA** | **NA** | **NA** | **NA** |
| 92 | *Hemidactylus mabouia* | Mabouia Clade | **NA** | **NA** | MW790327.1 | MW790458.1 | MW733817.1 |
| 93 | *Hemidactylus bowringii* | Tropical Asian | AF323515.1 | EU268405.1 | EU268374.1 | EU268313.1 | EU268343.1 |

| | | | | | | | |
|---|---|---|---|---|---|---|---|
| 94 | *Hemidactylus craspedotus* | Tropical Asian | **NA** | HM559586.1 | HM559618.1 | HM559684.1 | HM559651.1 |
| 95 | *Hemidactylus garnotii* | Tropical Asian | KY366207.1 | EU268396.1 | HM559631.1 | HM559697.1 | EU268333.1 |
| 96 | *Hemidactylus karenorum* | Tropical Asian | DQ120464.1 | EU268394.1 | **NA** | EU268301.1 | EU268331.1 |
| 97 | *Hemidactylus platyurus* | Tropical Asian | KY366206.1 | MN635567.1 | HM559619.1 | HM559685.1 | MW133495.1 |
| 98 | *Hemidactylus aquilonius* | Tropical Asian | KY366210.1 | **NA** | **NA** | KY366183.1 | **NA** |
| 99 | *Hemidactylus easai* | Tropical Asian | **NA** | **NA** | OM487119.1 | **NA** | **NA** |
| 100 | *Hemidactylus acanthopholis* | Tropical Asian | **NA** | MG711525.1 | MG711531.1 | MG711539.1 | MG711534.1 |
| 101 | *Hemidactylus aemulus* | Tropical Asian | **NA** | MZ274412.1 | **NA** | **NA** | **NA** |
| 102 | *Hemidactylus albofasciatus* | Tropical Asian | HM595679.1 | HM595643.1 | **NA** | KU720708.1 | **NA** |
| 103 | *Hemidactylus chikhaldaraensis* | Tropical Asian | **NA** | **NA** | MK569807.1 | **NA** | **NA** |
| 104 | *Hemidactylus chipkali* | Tropical Asian | **NA** | KX044190.2 | MK569809.1 | **NA** | **NA** |
| 105 | *Hemidactylus depressus* | Tropical Asian | **NA** | HM559589.1 | HM559621.1 | HM559687.1 | HM559654.1 |
| 106 | *Hemidactylus flavicaudus* | Tropical Asian | **NA** | **NA** | MN482234.1 | **NA** | **NA** |
| 107 | *Hemidactylus flaviviridis* | Tropical Asian | KY366202.1 | MN482349.1 | HM559628.1 | MN482264.1 | EU268324.1 |
| 108 | *Hemidactylus giganteus* | Tropical Asian | HM595694.1 | MZ274408.1 | MH454760.1 | MN482277.1 | HM622372.1 |
| 109 | *Hemidactylus gleadowi* | Tropical Asian | **NA** | MH454692.1 | MH454761.1 | MH454718.1 | MH454743.1 |
| 110 | *Hemidactylus gracilis* | Tropical Asian | HM595695.1 | MN482368.1 | MH454762.1 | MN482278.1 | MN482452.1 |
| 111 | *Hemidactylus graniticolus* | Tropical Asian | **NA** | MN482423.1 | MN482224.1 | MN482343.1 | MN482455.1 |
| 112 | *Hemidactylus gujaratensis* | Tropical Asian | **NA** | MG760350.1 | MG760344.1 | MG760347.1 | MG760351.1 |
| 113 | *Hemidactylus hegdei* | Tropical Asian | **NA** | OM685019.1 | **NA** | **NA** | **NA** |
| 114 | *Hemidactylus hemchandrai* | Tropical Asian | **NA** | MH454697.1 | MH454764.1 | MH454722.1 | MH454747.1 |
| 115 | *Hemidactylus hunae* | Tropical Asian | **NA** | HM559606.1 | HM559640.1 | HM559706.1 | HM559673.1 |
| 116 | *Hemidactylus kangerensis* | Tropical Asian | **NA** | KY938009.1 | MK569824.1 | **NA** | **NA** |
| 117 | *Hemidactylus kolliensis* | Tropical Asian | **NA** | **NA** | MK569825.1 | **NA** | **NA** |
| 118 | *Hemidactylus kushmorensis* | Tropical Asian | **NA** | MH454698.1 | **NA** | MH454724.1 | **NA** |
| 119 | *Hemidactylus lankae* | Tropical Asian | **NA** | MH454700.1 | HM559648.1 | MH454726.1 | MH454749.1 |
| 120 | *Hemidactylus leschenaultii* | Tropical Asian | HM595697.1 | MN482379.1 | MN482225.1 | MN482294.1 | MN482464.1 |
| 121 | *Hemidactylus mahonyi* | Tropical Asian | **NA** | **NA** | OM912578.1 | **NA** | **NA** |
| 122 | *Hemidactylus malcolmsmithii* | Tropical Asian | **NA** | **NA** | MK569842.1 | **NA** | **NA** |
| 123 | *Hemidactylus murrayi* | Tropical Asian | **NA** | MN482384.1 | **NA** | MN482298.1 | **NA** |
| 124 | *Hemidactylus paaragowli* | Tropical Asian | **NA** | MK852285.1 | MK802893.1 | MK852288.1 | MK852280.1 |

| | | | | | | | |
|---|---|---|---|---|---|---|---|
| 125 | *Hemidactylus pakkamalaiensis* | Tropical Asian | **NA** | **NA** | MN482224.1 | **NA** | **NA** |
| 126 | *Hemidactylus parvimaculatus* | Tropical Asian | **NA** | MH454704.1 | MH454766.1 | MH454730.1 | MH454754.1 |
| 127 | *Hemidactylus raya* | Tropical Asian | **NA** | MZ274414.1 | **NA** | **NA** | **NA** |
| 128 | *Hemidactylus reticulatus* | Tropical Asian | HM595707.1 | MN482392.1 | MH454767.1 | MN482306.1 | MN482476.1 |
| 129 | *Hemidactylus rishivalleyensis* | Tropical Asian | **NA** | **NA** | MT773224.1 | **NA** | **NA** |
| 130 | *Hemidactylus sahgali* | Tropical Asian | **NA** | MN482393.1 | MN482229.1 | MN482310.1 | MN482479.1 |
| 131 | *Hemidactylus sankariensis* | Tropical Asian | **NA** | **NA** | MK569844.1 | **NA** | **NA** |
| 132 | *Hemidactylus sataraensis* | Tropical Asian | HM595708.1 | MH454706.1 | MH454768.1 | MH454732.1 | MH454756.1 |
| 133 | *Hemidactylus saxicolus* | Tropical Asian | **NA** | MZ274419.1 | **NA** | **NA** | **NA** |
| 134 | *Hemidactylus scabriceps* | Tropical Asian | KX902974.1 | KX902975.1 | MH454769.1 | MH454733.1 | KX902976.1 |
| 135 | *Hemidactylus sirumalaiensis* | Tropical Asian | **NA** | **NA** | MT943051.1 | **NA** | **NA** |
| 136 | *Hemidactylus siva* | Tropical Asian | **NA** | MG438452.1 | MK569845.1 | MG438451.1 | MG438453.1 |
| 137 | *Hemidactylus srikanthani* | Tropical Asian | **NA** | **NA** | OM912580.1 | **NA** | **NA** |
| 138 | *Hemidactylus sushilduttai* | Tropical Asian | **NA** | MN482411.1 | MN482236.1 | MN482341.1 | MN482495.1 |
| 139 | *Hemidactylus tamhiniensis* | Tropical Asian | **NA** | **NA** | MN482222.1 | **NA** | **NA** |
| 140 | *Hemidactylus treutleri* | Tropical Asian | **NA** | MN482422.1 | MN482237.1 | MN482320.1 | MN482506.1 |
| 141 | *Hemidactylus vanam* | Tropical Asian | **NA** | MG711529.1 | MN482239.1 | MG711542.1 | MG711535.1 |
| 142 | *Hemidactylus varadgirii* | Tropical Asian | **NA** | MN482424.1 | **NA** | MN482344.1 | MN482507.1 |
| 143 | *Hemidactylus vijayraghavani* | Tropical Asian | **NA** | MN482426.1 | **NA** | MN482322.1 | MN482509.1 |
| 144 | *Hemidactylus whitakeri* | Tropical Asian | **NA** | MN482429.1 | **NA** | MN482322.1 | MN482512.1 |
| 145 | *Hemidactylus xericolus* | Tropical Asian | **NA** | **NA** | MN482235.1 | **NA** | **NA** |
| 146 | *Hemidactylus yajurvedi* | Tropical Asian | **NA** | KT601565.1 | MH454772.1 | KT601568.1 | KT601567.1 |
| 147 | *Hemidactylus frenatus* | Tropical Asian | KY366200.1 | KT455028.1 | HM559630.1 | HM559695.1 | EU268328.1 |
| 148 | *Hemidactylus aaronbaueri* | Tropical Asian | HM595677.1 | HM595640.1 | MN482222.1 | HM622352.1 | HM622367.1 |
| 149 | *Hemidactylus maculatus* | Tropical Asian | HM595700.1 | MN482382.1 | MN482227.1 | MN482297.1 | MN482466.1 |
| 150 | *Hemidactylus prashadi* | Tropical Asian | **NA** | MN482386.1 | MN482228.1 | MN482301.1 | MN482469.1 |
| 151 | *Hemidactylus triedrus* | Tropical Asian | HM595711.1 | MG742361.1 | MH666065.1 | MH454735.1 | MH454759.1 |
| 152 | *Hemidactylus tenkatei* | Tropical Asian | **NA** | **NA** | KM975943.1 | KM881691.1 | **NA** |
