## Supplementary figures and images for "Globetrotting geckos: Historical biogeography suggests an Indian origin and “Out-Of-India” dispersal for the cosmopolitan *Hemidactylus* geckos"

### BioGeoBEARS 5 area result

**Hemidactylus DEC with distdisppadj**

**ancstates: global optim, 2 areas max. d=0.0061; e=0.0041; j=0; LnL=−127.94**

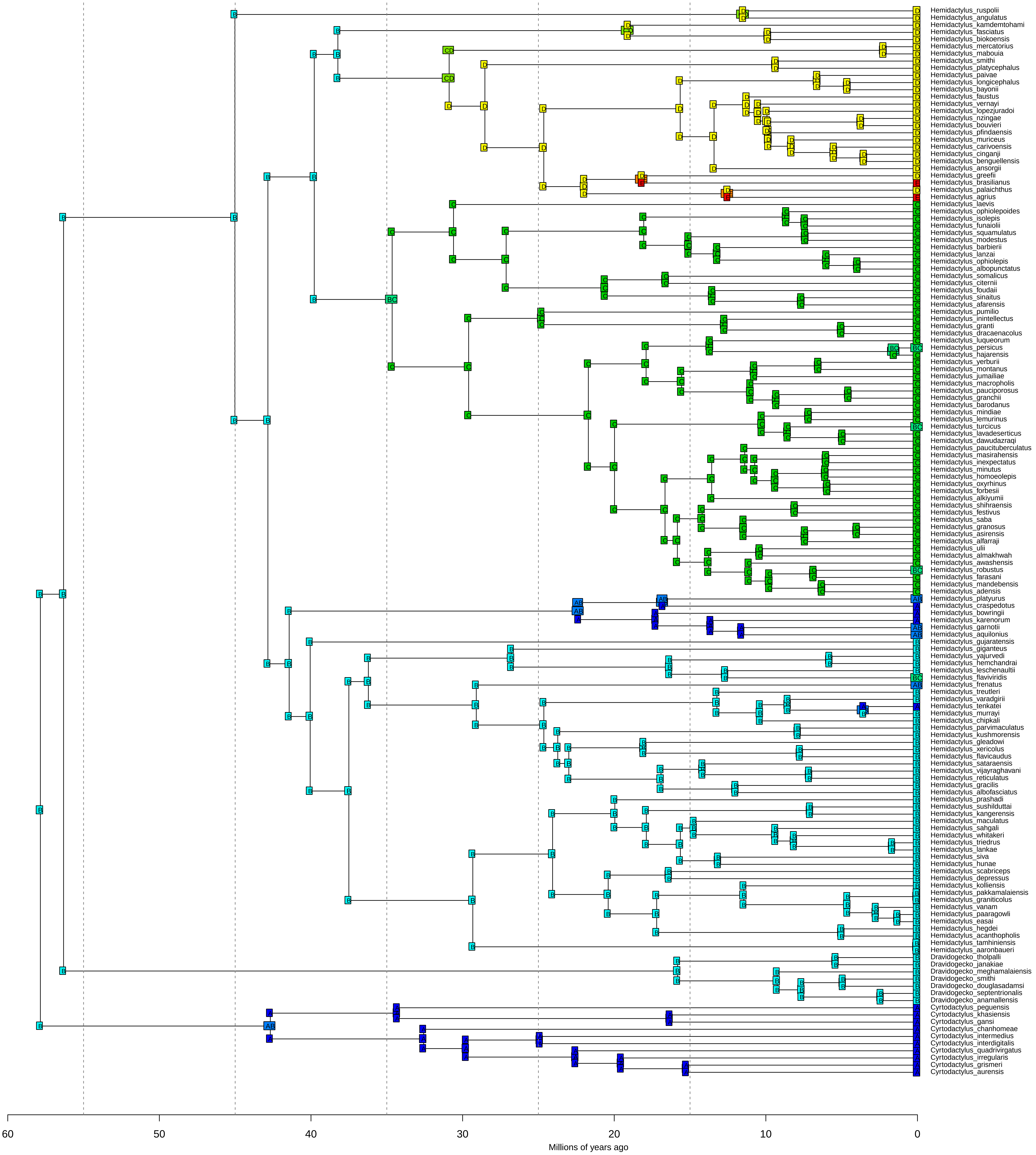

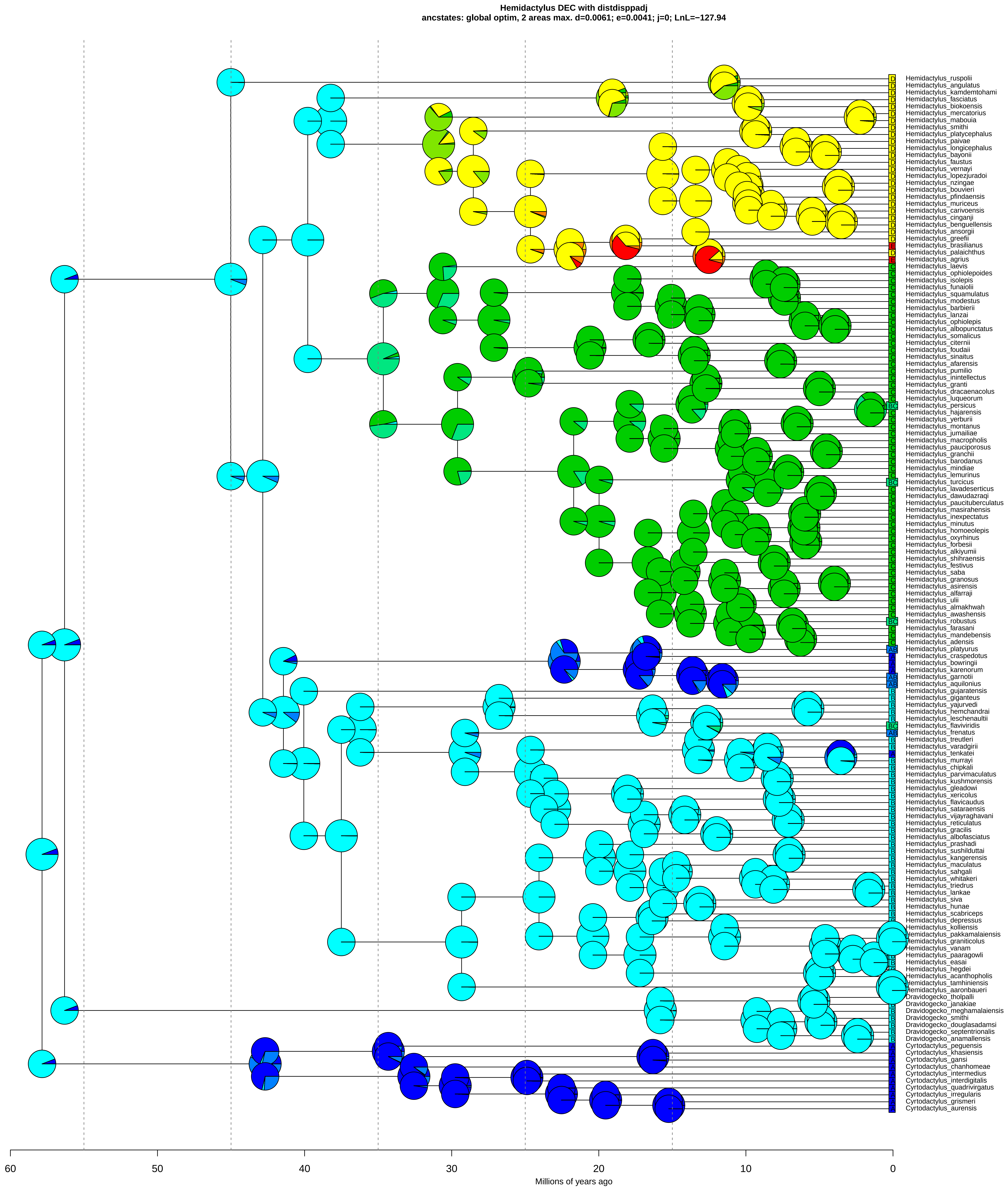

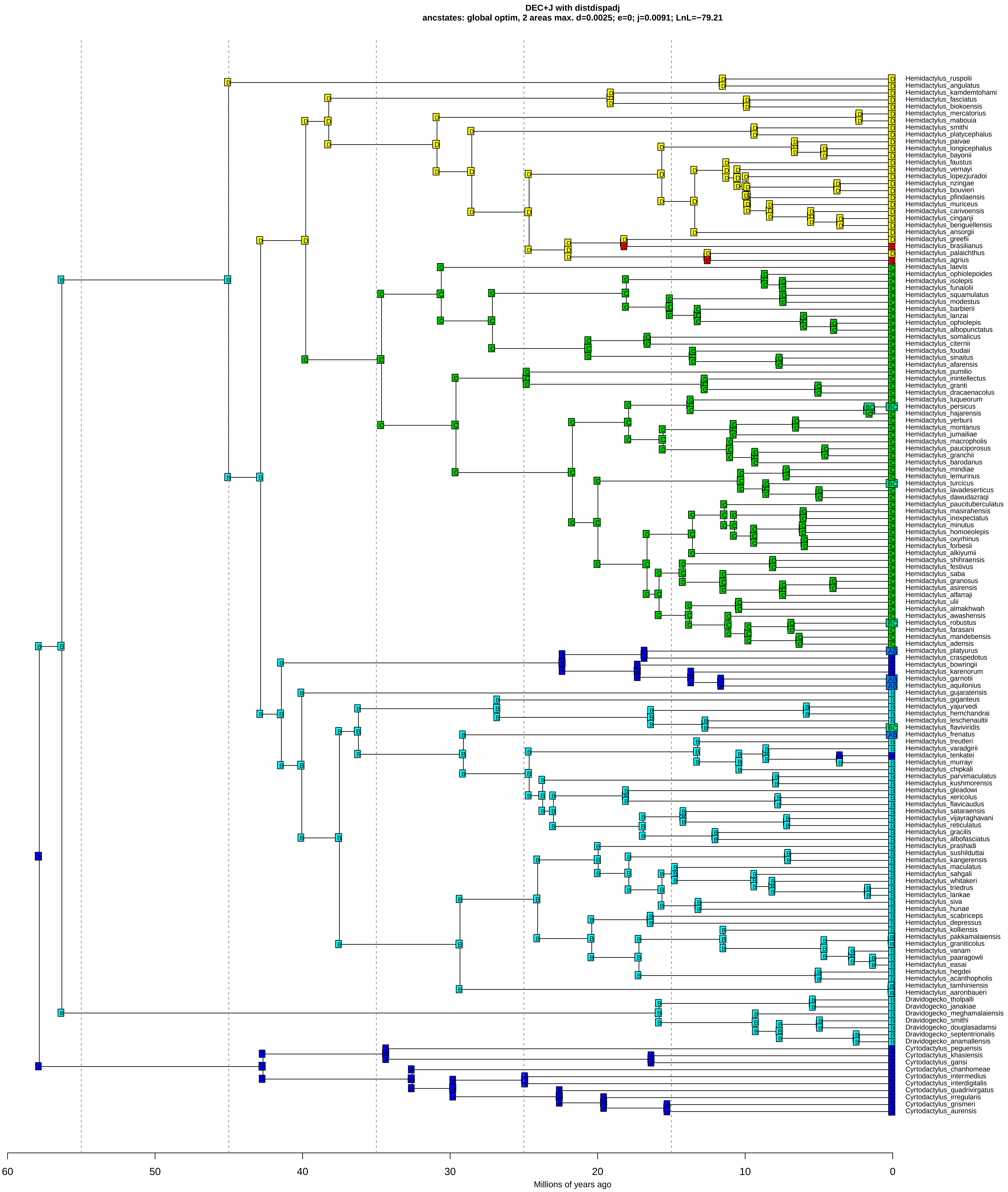

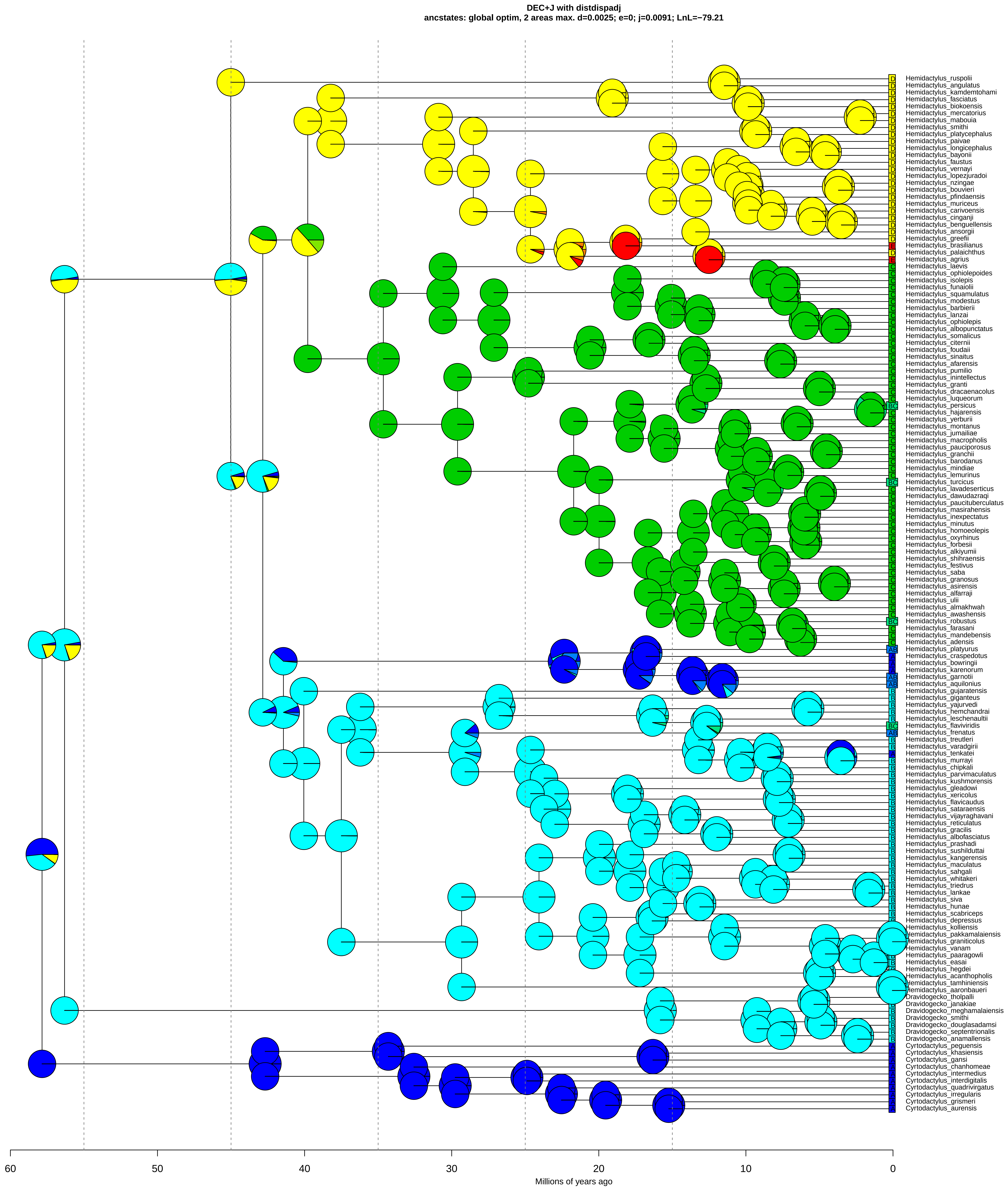

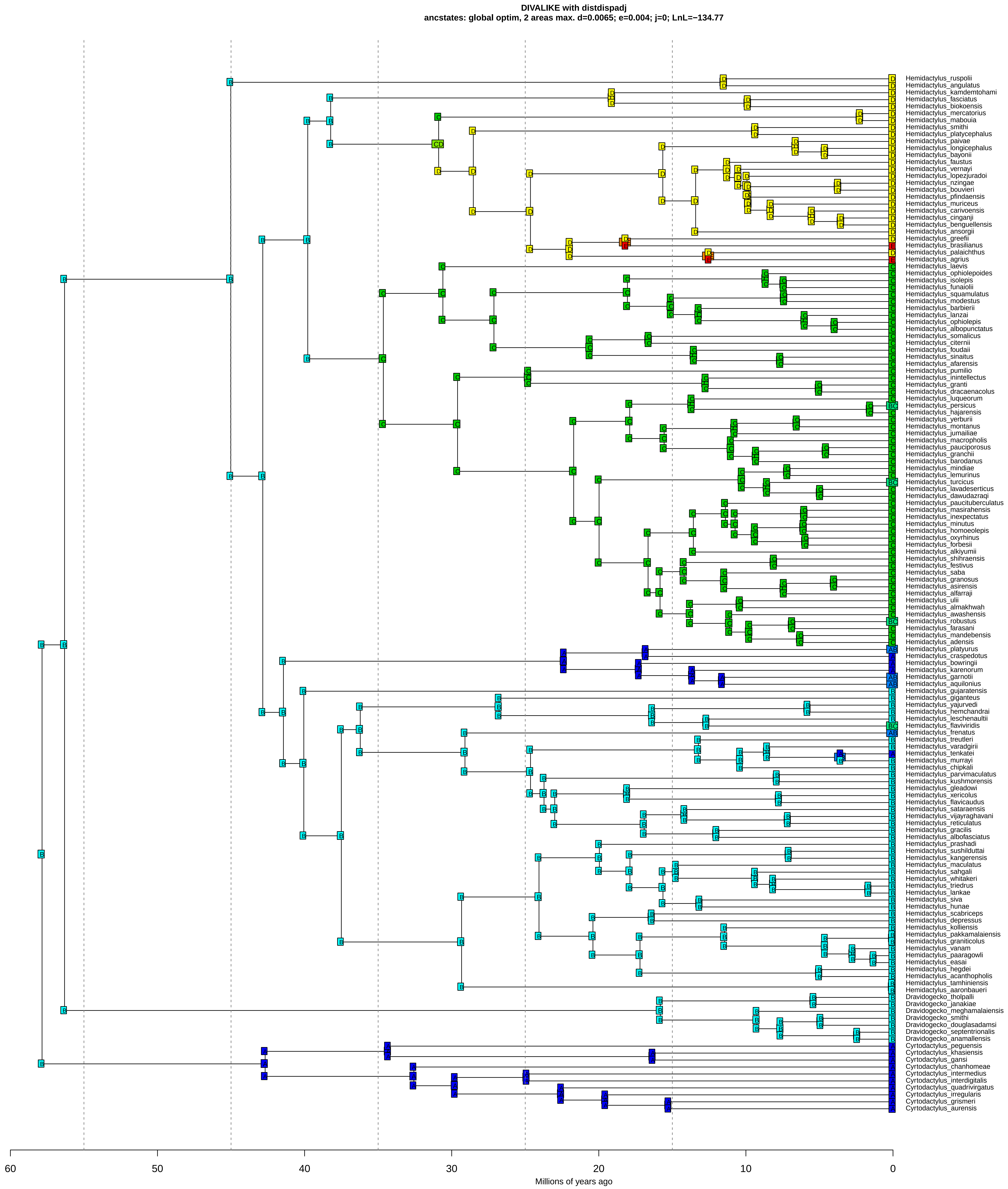

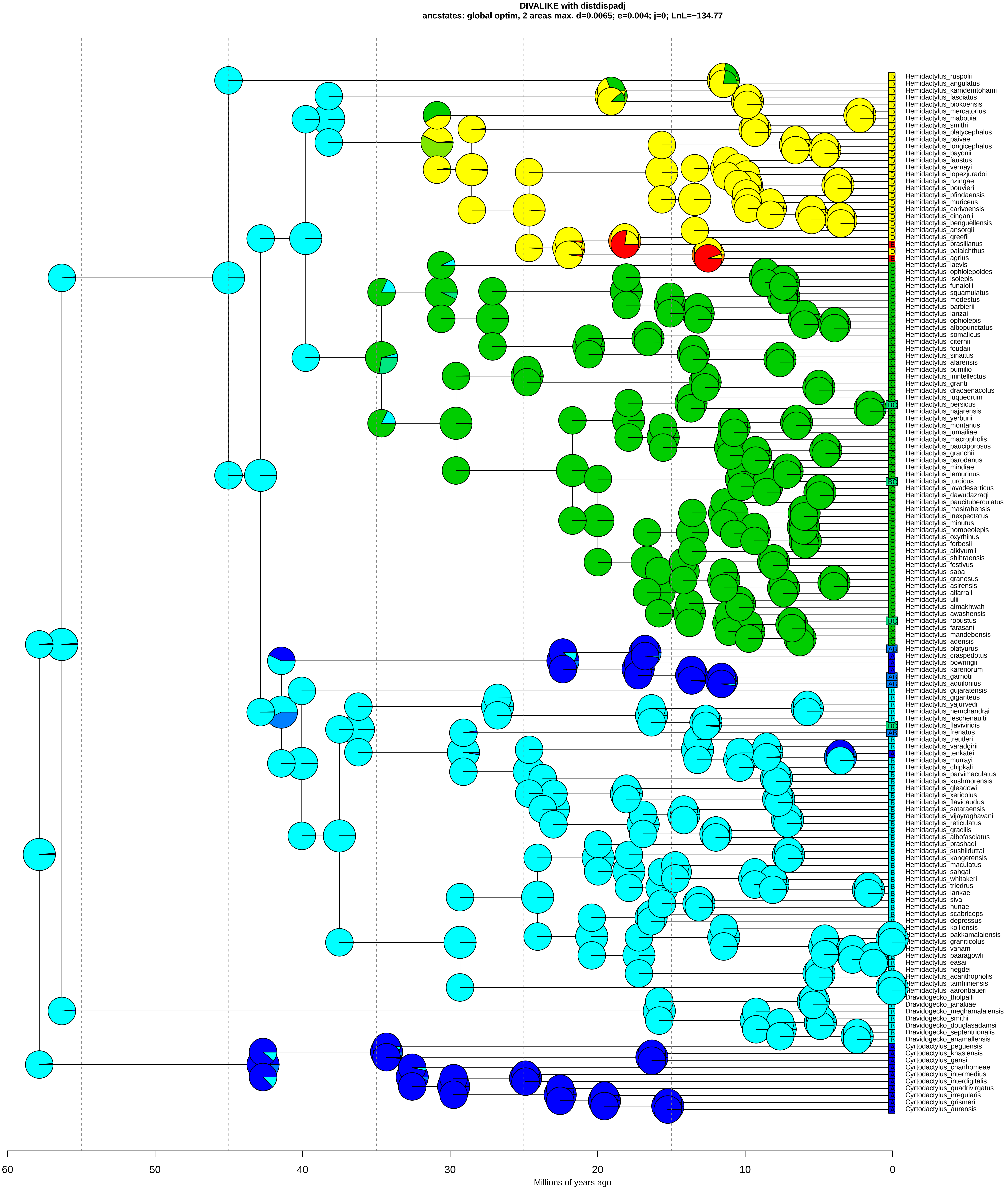

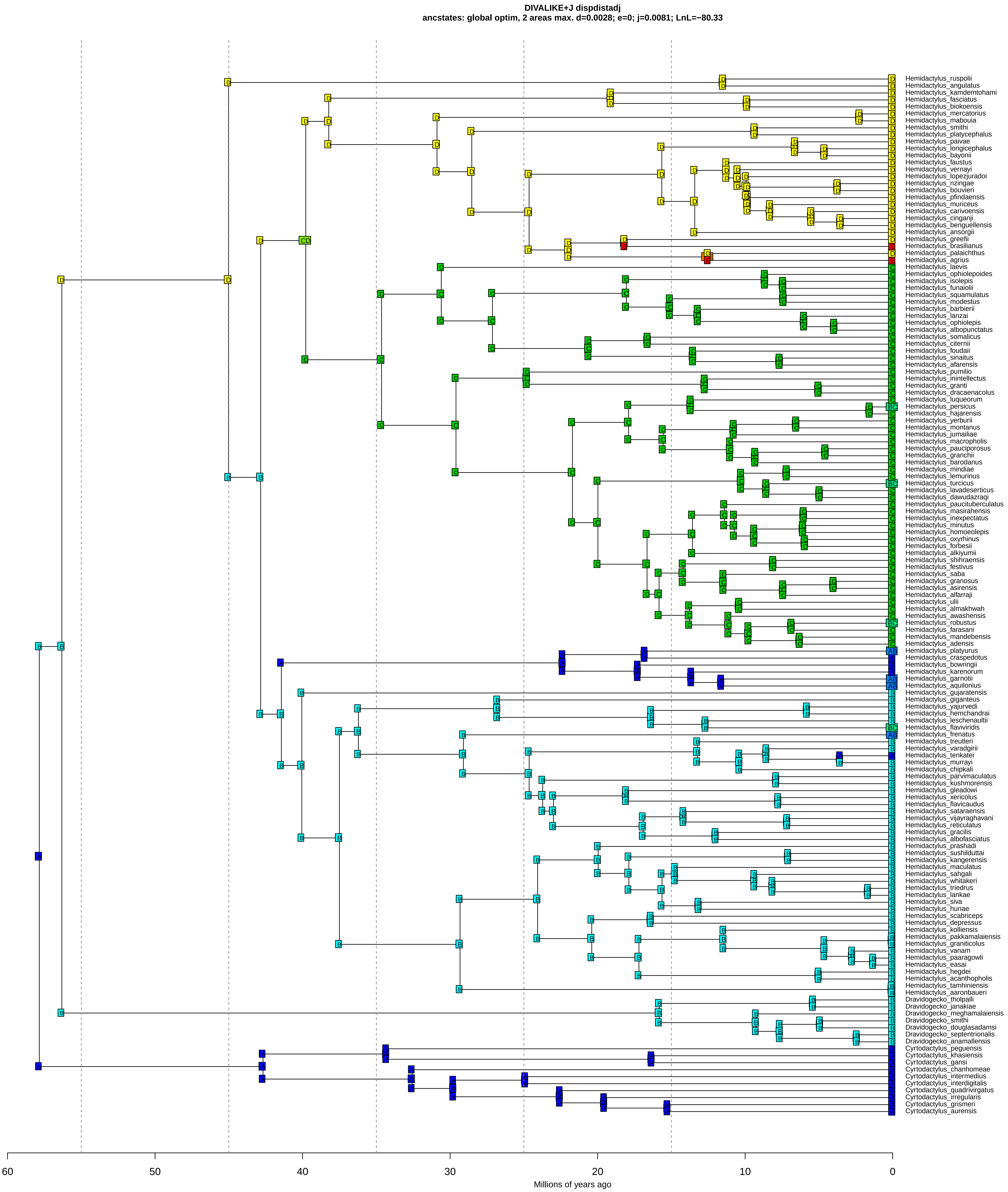

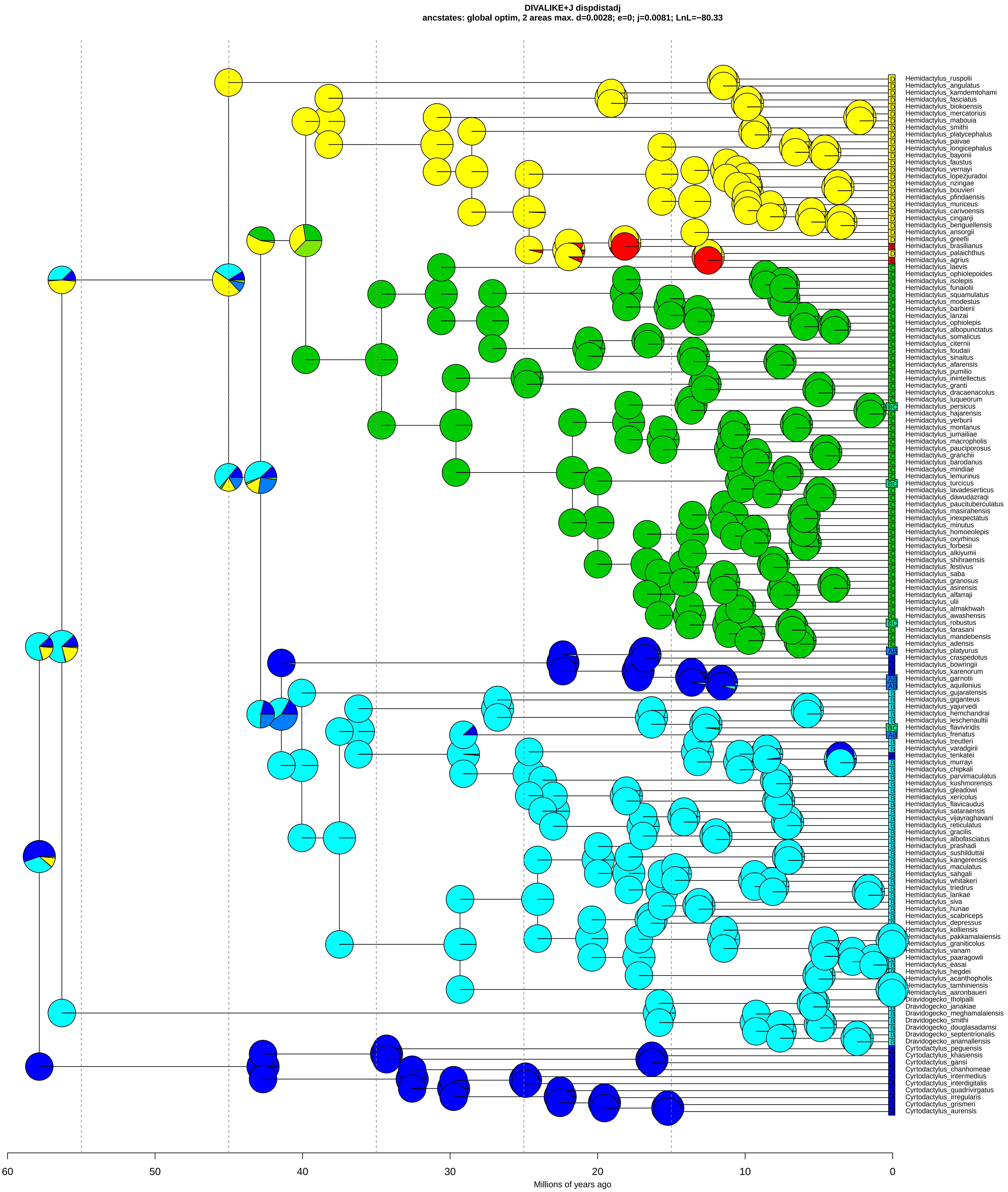

### Coalescent phylogeny

**Supplementary Figure 1 (S1)**

Species tree constructed using StarBeast

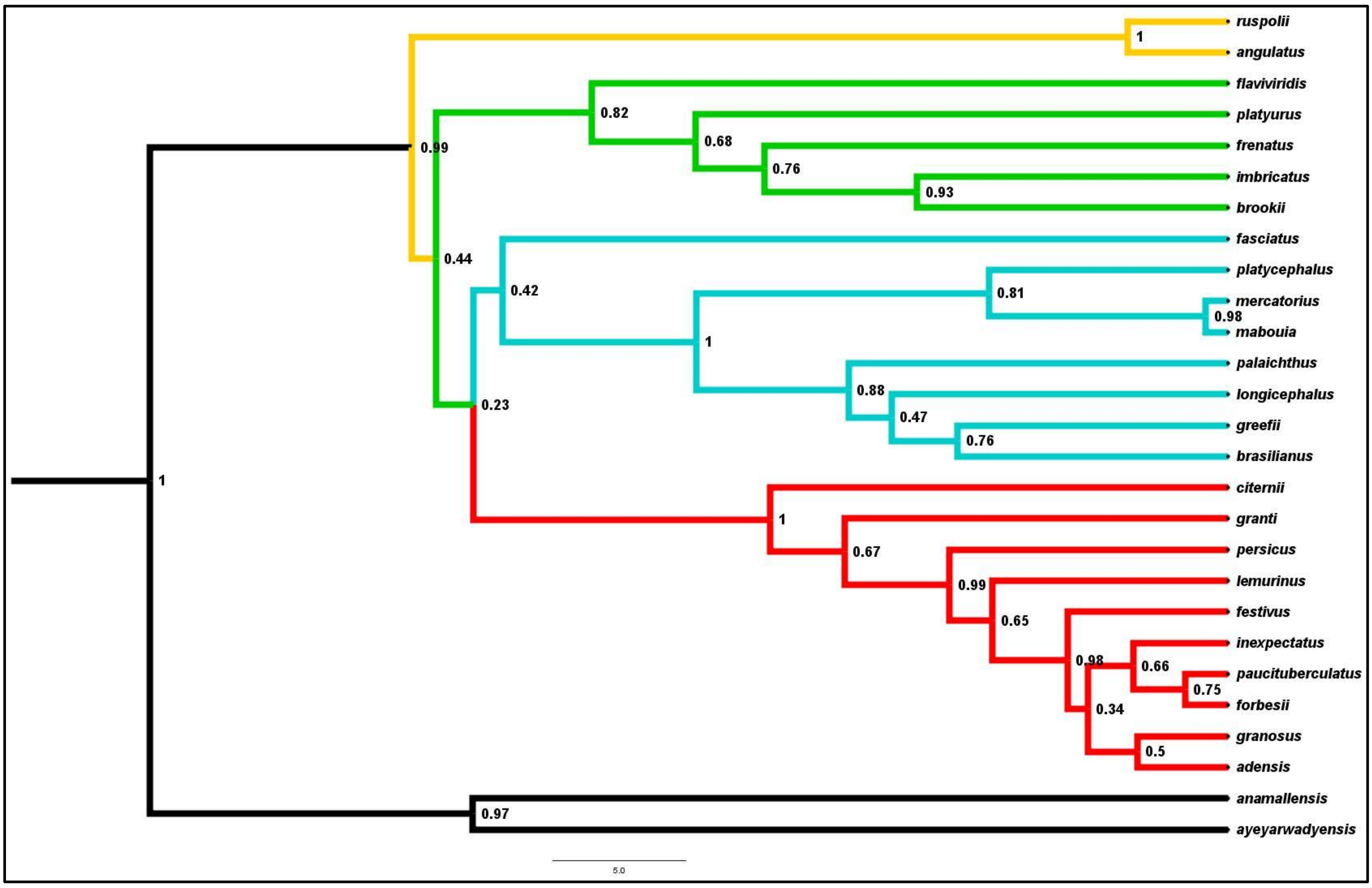
