## Supplementary material for "Globetrotting geckos: Historical biogeography suggests an Indian origin and “Out-Of-India” dispersal for the cosmopolitan *Hemidactylus* geckos": BioGeoBEARS result without sister groups

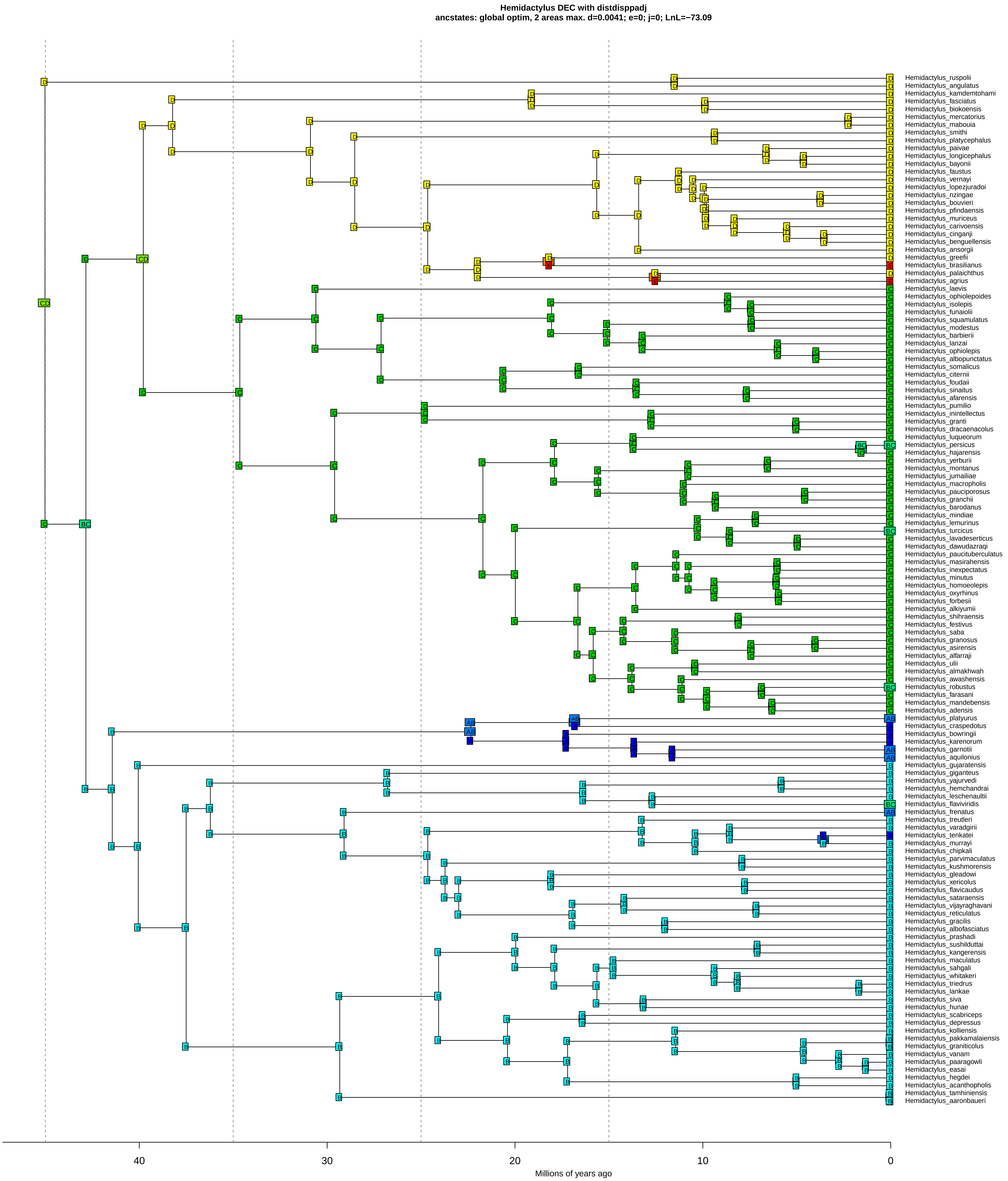

Hemidactylus DEC with distdisppadj
ancstates: global optim, 2 areas max. d=0.0041; e=0; j=0; LnL=−73.09
Hemidactylus_ruspolii
Hemidactylus_angulatus
Hemidactylus_kamdemtohami
Hemidactylus_fasciatus
Hemidactylus_biokoensis
Hemidactylus_mercatorius
Hemidactylus_mabouia
Hemidactylus_smithi
Hemidactylus_platycephalus
Hemidactylus_paivae
Hemidactylus_longicephalus
Hemidactylus_bayonii
Hemidactylus_faustus
Hemidactylus_vernayi
Hemidactylus_lopezjuradoi
Hemidactylus_nzingae
Hemidactylus_bouvieri
Hemidactylus_pfindaensis
Hemidactylus_muriceus
Hemidactylus_carivoensis
Hemidactylus_cinganji
Hemidactylus_benguellensis
Hemidactylus_ansorgii
Hemidactylus_greefii
Hemidactylus_brasilianus
Hemidactylus_palaichthus
Hemidactylus_agrius
Hemidactylus_laevis
Hemidactylus_ophiolepoides
Hemidactylus_isolepis
Hemidactylus_funaiolii
Hemidactylus_squamulatus
Hemidactylus_modestus
Hemidactylus_barbierii
Hemidactylus_lanzai
Hemidactylus_ophiolepis
Hemidactylus_albopunctatus
Hemidactylus_somalicus
Hemidactylus_citernii
Hemidactylus_foudaii
Hemidactylus_sinaitus
Hemidactylus_afarensis
Hemidactylus_pumilio
Hemidactylus_inintellectus
Hemidactylus_granti
Hemidactylus_dracaenacolus
Hemidactylus_luqueorum
Hemidactylus_persicus
Hemidactylus_hajarensis
Hemidactylus_yerburii
Hemidactylus_montanus
Hemidactylus_jumailiae
Hemidactylus_macropholis
Hemidactylus_pauciporosus
Hemidactylus_granchii
Hemidactylus_barodanus
Hemidactylus_mindiae
Hemidactylus_lemurinus
Hemidactylus_turcicus
Hemidactylus_lavadeserticus
Hemidactylus_dawudazraqi
Hemidactylus_paucituberculatus
Hemidactylus_masirahensis
Hemidactylus_inexpectatus
Hemidactylus_minutus
Hemidactylus_homoeolepis
Hemidactylus_oxyrhinus
Hemidactylus_forbesii
Hemidactylus_alkiyumii
Hemidactylus_shihraensis
Hemidactylus_festivus
Hemidactylus_saba
Hemidactylus_granosus
Hemidactylus_asirensis
Hemidactylus_alfarraji
Hemidactylus_ulii
Hemidactylus_almakhwah
Hemidactylus_awashensis
Hemidactylus_robustus
Hemidactylus_farasani
Hemidactylus_mandebensis
Hemidactylus_adensis
Hemidactylus_platyurus
Hemidactylus_craspedotus
Hemidactylus_bowringii
Hemidactylus_karenorum
Hemidactylus_garnotii
Hemidactylus_aquilonius
Hemidactylus_gujaratensis
Hemidactylus_giganteus
Hemidactylus_yajurvedi
Hemidactylus_hemchandrai
Hemidactylus_leschenaultii
Hemidactylus_flaviviridis
Hemidactylus_frenatus
Hemidactylus_treutleri
Hemidactylus_varadgirii
Hemidactylus_tenkatei
Hemidactylus_murrayi
Hemidactylus_chipkali
Hemidactylus_parvimaculatus
Hemidactylus_kushmorensis
Hemidactylus_gleadowi
Hemidactylus_xericolus
Hemidactylus_flavicaudus
Hemidactylus_sataraensis
Hemidactylus_vijayraghavani
Hemidactylus_reticulatus
Hemidactylus_gracilis
Hemidactylus_albofasciatus
Hemidactylus_prashadi
Hemidactylus_sushilduttai
Hemidactylus_kangerensis
Hemidactylus_maculatus
Hemidactylus_sahgali
Hemidactylus_whitakeri
Hemidactylus_triedrus
Hemidactylus_lankae
Hemidactylus_siva
Hemidactylus_hunae
Hemidactylus_scabriceps
Hemidactylus_depressus
Hemidactylus_kolliensis
Hemidactylus_pakkamalaiensis
Hemidactylus_graniticolus
Hemidactylus_vanam
Hemidactylus_paaragowli
Hemidactylus_easai
Hemidactylus_hegdei
Hemidactylus_acanthopholis
Hemidactylus_tamhiniensis
Hemidactylus_aaronbaueri
40
30
20
10
0
Millions of years ago

**Hemidactylus DEC with distdisppadj**
**ancstates: global optim, 2 areas max. d=0.0041; e=0; j=0; LnL=−73.09**

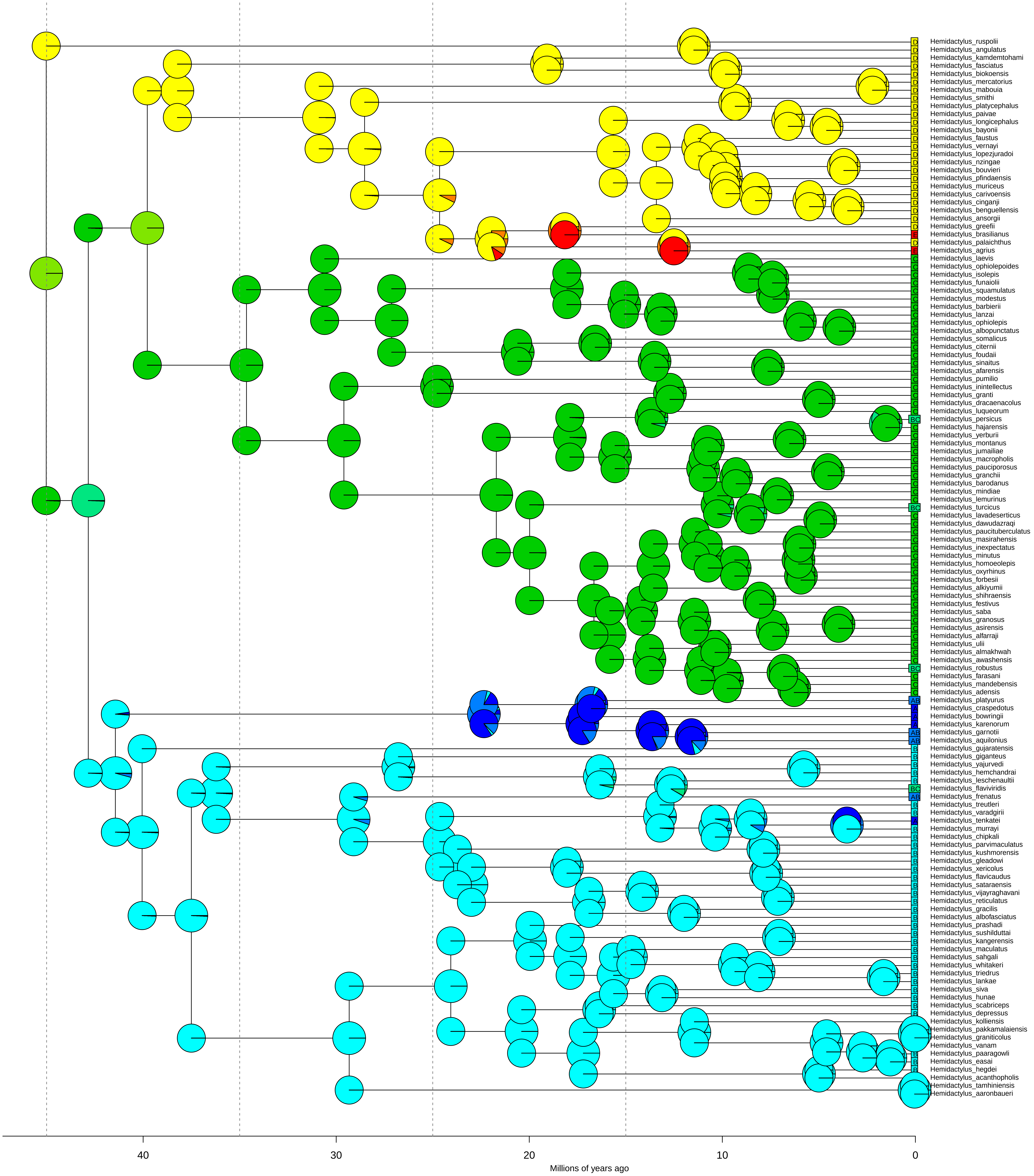

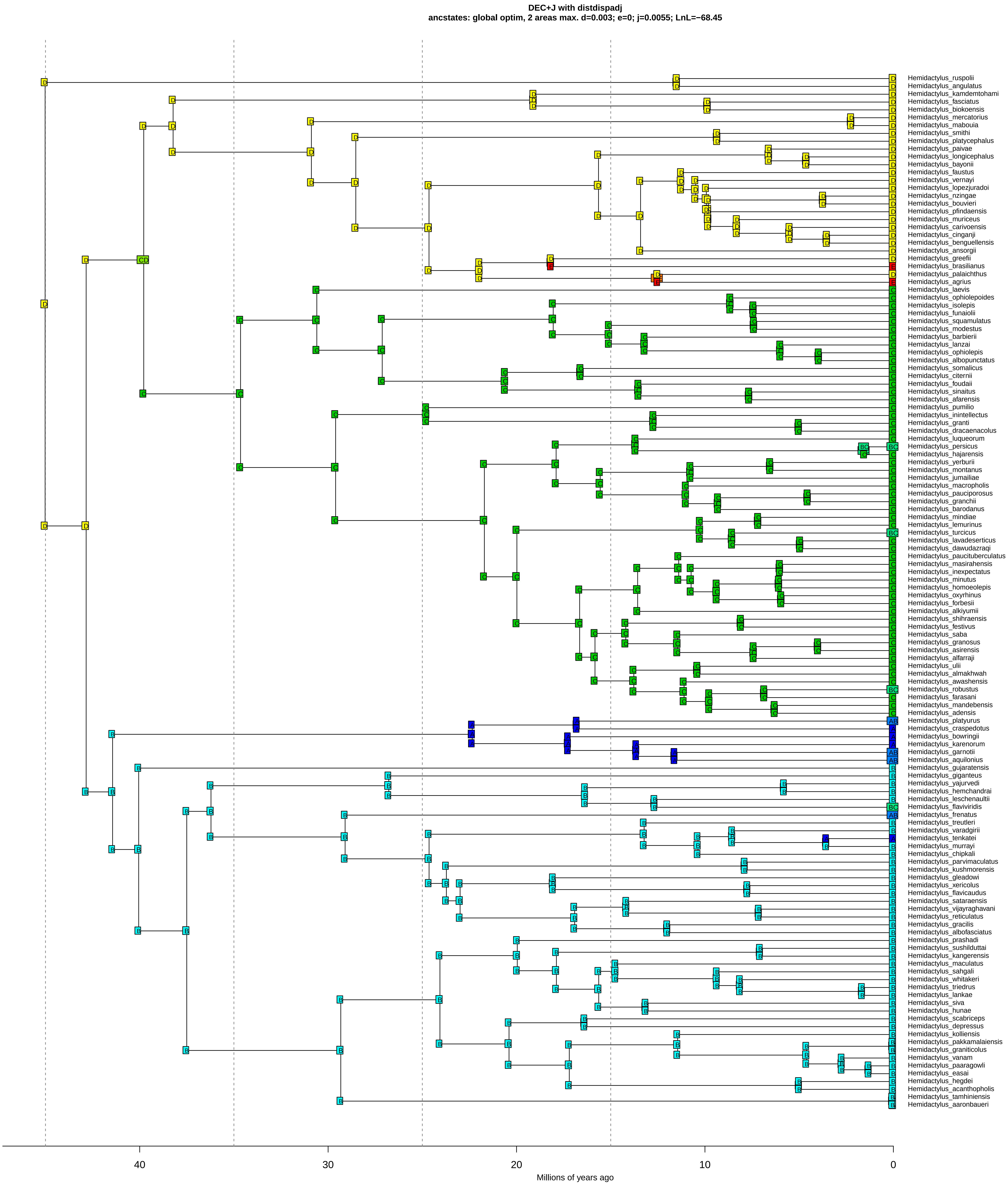

DEC+J with distdispadj
ancstates: global optim, 2 areas max. d=0.003; e=0; j=0.0055; LnL=−68.45
Hemidactylus_ruspolii
Hemidactylus_angulatus
Hemidactylus_kamdemtohami
Hemidactylus_fasciatus
Hemidactylus_biokoensis
Hemidactylus_mercatorius
Hemidactylus_mabouia
Hemidactylus_smithi
Hemidactylus_platycephalus
Hemidactylus_paivae
Hemidactylus_longicephalus
Hemidactylus_bayonii
Hemidactylus_faustus
Hemidactylus_vernayi
Hemidactylus_lopezjuradoi
Hemidactylus_nzingae
Hemidactylus_bouvieri
Hemidactylus_pfindaensis
Hemidactylus_muriceus
Hemidactylus_carivoensis
Hemidactylus_cinganji
Hemidactylus_benguellensis
Hemidactylus_ansorgii
Hemidactylus_greefii
Hemidactylus_brasilianus
Hemidactylus_palaichthus
Hemidactylus_agrius
Hemidactylus_laevis
Hemidactylus_ophiolepoides
Hemidactylus_isolepis
Hemidactylus_funaiolii
Hemidactylus_squamulatus
Hemidactylus_modestus
Hemidactylus_barbierii
Hemidactylus_lanzai
Hemidactylus_ophiolepis
Hemidactylus_albopunctatus
Hemidactylus_somalicus
Hemidactylus_citernii
Hemidactylus_foudaii
Hemidactylus_sinaitus
Hemidactylus_afarensis
Hemidactylus_pumilio
Hemidactylus_inintellectus
Hemidactylus_granti
Hemidactylus_dracaenacolus
Hemidactylus_luqueorum
Hemidactylus_persicus
Hemidactylus_hajarensis
Hemidactylus_yerburii
Hemidactylus_montanus
Hemidactylus_jumailiae
Hemidactylus_macropholis
Hemidactylus_pauciporosus
Hemidactylus_granchii
Hemidactylus_barodanus
Hemidactylus_mindiae
Hemidactylus_lemurinus
Hemidactylus_turcicus
Hemidactylus_lavadeserticus
Hemidactylus_dawudazraqi
Hemidactylus_paucituberculatus
Hemidactylus_masirahensis
Hemidactylus_inexpectatus
Hemidactylus_minutus
Hemidactylus_homoeolepis
Hemidactylus_oxyrhinus
Hemidactylus_forbesii
Hemidactylus_alkiyumii
Hemidactylus_shihraensis
Hemidactylus_festivus
Hemidactylus_saba
Hemidactylus_granosus
Hemidactylus_asirensis
Hemidactylus_alfarraji
Hemidactylus_ulii
Hemidactylus_almakhwah
Hemidactylus_awashensis
Hemidactylus_robustus
Hemidactylus_farasani
Hemidactylus_mandebensis
Hemidactylus_adensis
Hemidactylus_platyurus
Hemidactylus_craspedotus
Hemidactylus_bowringii
Hemidactylus_karenorum
Hemidactylus_garnotii
Hemidactylus_aquilonius
Hemidactylus_gujaratensis
Hemidactylus_giganteus
Hemidactylus_yajurvedi
Hemidactylus_hemchandrai
Hemidactylus_leschenaultii
Hemidactylus_flaviviridis
Hemidactylus_frenatus
Hemidactylus_treutleri
Hemidactylus_varadgirii
Hemidactylus_tenkatei
Hemidactylus_murrayi
Hemidactylus_chipkali
Hemidactylus_parvimaculatus
Hemidactylus_kushmorensis
Hemidactylus_gleadowi
Hemidactylus_xericolus
Hemidactylus_flavicaudus
Hemidactylus_sataraensis
Hemidactylus_vijayraghavani
Hemidactylus_reticulatus
Hemidactylus_gracilis
Hemidactylus_albofasciatus
Hemidactylus_prashadi
Hemidactylus_sushilduttai
Hemidactylus_kangerensis
Hemidactylus_maculatus
Hemidactylus_sahgali
Hemidactylus_whitakeri
Hemidactylus_triedrus
Hemidactylus_lankae
Hemidactylus_siva
Hemidactylus_hunae
Hemidactylus_scabriceps
Hemidactylus_depressus
Hemidactylus_kolliensis
Hemidactylus_pakkamalaiensis
Hemidactylus_graniticolus
Hemidactylus_vanam
Hemidactylus_paaragowli
Hemidactylus_easai
Hemidactylus_hegdei
Hemidactylus_acanthopholis
Hemidactylus_tamhiniensis
Hemidactylus_aaronbaueri
40
30
20
10
0
Millions of years ago

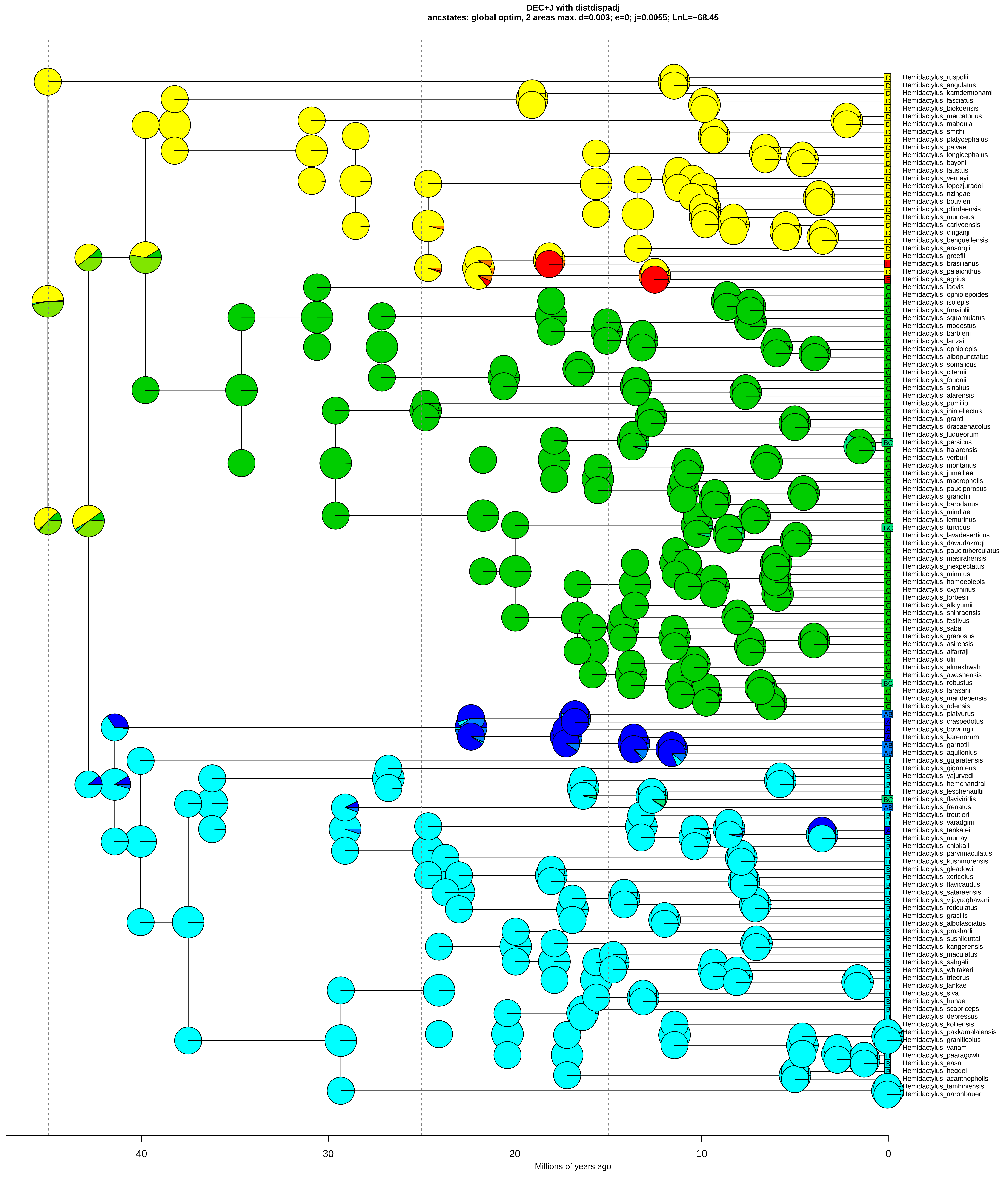

DEC+J with distdispadj
ancstates: global optim, 2 areas max. d=0.003; e=0; j=0.0055; LnL=−68.45
Hemidactylus_ruspolii
Hemidactylus_angulatus
Hemidactylus_kamdemtohami
Hemidactylus_fasciatus
Hemidactylus_biokoensis
Hemidactylus_mercatorius
Hemidactylus_mabouia
Hemidactylus_smithi
Hemidactylus_platycephalus
Hemidactylus_paivae
Hemidactylus_longicephalus
Hemidactylus_bayonii
Hemidactylus_faustus
Hemidactylus_vernayi
Hemidactylus_lopezjuradoi
Hemidactylus_nzingae
Hemidactylus_bouvieri
Hemidactylus_pfindaensis
Hemidactylus_muriceus
Hemidactylus_carivoensis
Hemidactylus_cinganji
Hemidactylus_benguellensis
Hemidactylus_ansorgii
Hemidactylus_greefii
Hemidactylus_brasilianus
Hemidactylus_palaichthus
Hemidactylus_agrius
Hemidactylus_laevis
Hemidactylus_ophiolepoides
Hemidactylus_isolepis
Hemidactylus_funaiolii
Hemidactylus_squamulatus
Hemidactylus_modestus
Hemidactylus_barbierii
Hemidactylus_lanzai
Hemidactylus_ophiolepis
Hemidactylus_albopunctatus
Hemidactylus_somalicus
Hemidactylus_citernii
Hemidactylus_foudaii
Hemidactylus_sinaitus
Hemidactylus_afarensis
Hemidactylus_pumilio
Hemidactylus_inintellectus
Hemidactylus_granti
Hemidactylus_dracaenacolus
Hemidactylus_luqueorum
Hemidactylus_persicus
Hemidactylus_hajarensis
Hemidactylus_yerburii
Hemidactylus_montanus
Hemidactylus_jumailiae
Hemidactylus_macropholis
Hemidactylus_pauciporosus
Hemidactylus_granchii
Hemidactylus_barodanus
Hemidactylus_mindiae
Hemidactylus_lemurinus
Hemidactylus_turcicus
Hemidactylus_lavadeserticus
Hemidactylus_dawudazraqi
Hemidactylus_paucituberculatus
Hemidactylus_masirahensis
Hemidactylus_inexpectatus
Hemidactylus_minutus
Hemidactylus_homoeolepis
Hemidactylus_oxyrhinus
Hemidactylus_forbesii
Hemidactylus_alkiyumii
Hemidactylus_shihraensis
Hemidactylus_festivus
Hemidactylus_saba
Hemidactylus_granosus
Hemidactylus_asirensis
Hemidactylus_alfarraji
Hemidactylus_ulii
Hemidactylus_almakhwah
Hemidactylus_awashensis
Hemidactylus_robustus
Hemidactylus_farasani
Hemidactylus_mandebensis
Hemidactylus_adensis
Hemidactylus_platyurus
Hemidactylus_craspedotus
Hemidactylus_bowringii
Hemidactylus_karenorum
Hemidactylus_garnotii
Hemidactylus_aquilonius
Hemidactylus_gujaratensis
Hemidactylus_giganteus
Hemidactylus_yajurvedi
Hemidactylus_hemchandrai
Hemidactylus_leschenaultii
Hemidactylus_flaviviridis
Hemidactylus_frenatus
Hemidactylus_treutleri
Hemidactylus_varadgirii
Hemidactylus_tenkatei
Hemidactylus_murrayi
Hemidactylus_chipkali
Hemidactylus_parvimaculatus
Hemidactylus_kushmorensis
Hemidactylus_gleadowi
Hemidactylus_xericolus
Hemidactylus_flavicaudus
Hemidactylus_sataraensis
Hemidactylus_vijayraghavani
Hemidactylus_reticulatus
Hemidactylus_gracilis
Hemidactylus_albofasciatus
Hemidactylus_prashadi
Hemidactylus_sushilduttai
Hemidactylus_kangerensis
Hemidactylus_maculatus
Hemidactylus_sahgali
Hemidactylus_whitakeri
Hemidactylus_triedrus
Hemidactylus_lankae
Hemidactylus_siva
Hemidactylus_hunae
Hemidactylus_scabriceps
Hemidactylus_depressus
Hemidactylus_kolliensis
Hemidactylus_pakkamalaiensis
Hemidactylus_graniticolus
Hemidactylus_vanam
Hemidactylus_paaragowli
Hemidactylus_easai
Hemidactylus_hegdei
Hemidactylus_acanthopholis
Hemidactylus_tamhiniensis
Hemidactylus_aaronbaueri
40
30
20
10
0
Millions of years ago

**DIVALIKE with distdispadj**
**ancstates: global optim, 2 areas max. d=0.0044; e=0; j=0; LnL=−70.83**

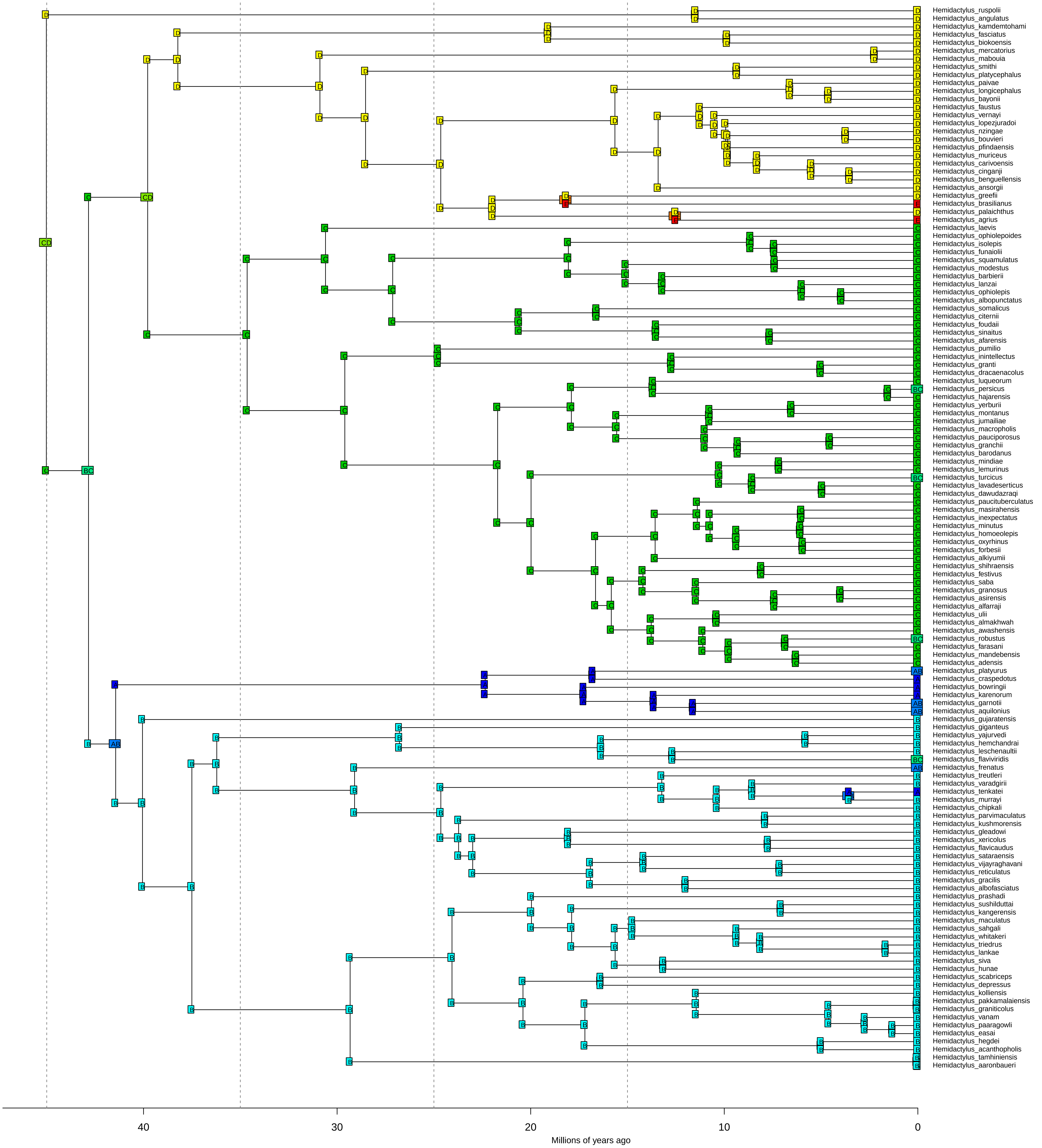

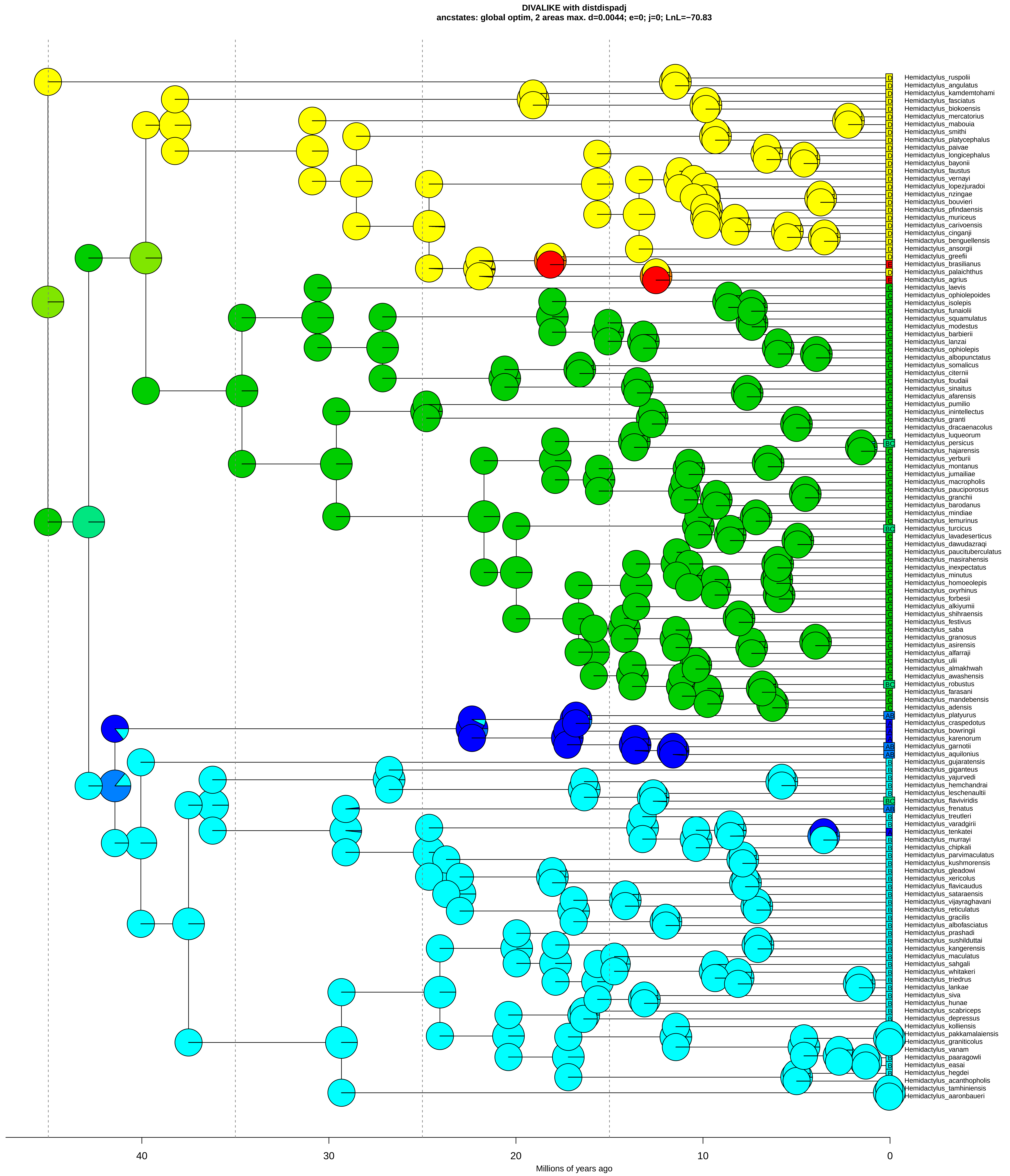

DIVALIKE with distdispadj
ancstates: global optim, 2 areas max. d=0.0044; e=0; j=0; LnL=−70.83
Hemidactylus_ruspolii
Hemidactylus_angulatus
Hemidactylus_kamdemtohami
Hemidactylus_fasciatus
Hemidactylus_biokoensis
Hemidactylus_mercatorius
Hemidactylus_mabouia
Hemidactylus_smithi
Hemidactylus_platycephalus
Hemidactylus_paivae
Hemidactylus_longicephalus
Hemidactylus_bayonii
Hemidactylus_faustus
Hemidactylus_vernayi
Hemidactylus_lopezjuradoi
Hemidactylus_nzingae
Hemidactylus_bouvieri
Hemidactylus_pfindaensis
Hemidactylus_muriceus
Hemidactylus_carivoensis
Hemidactylus_cinganji
Hemidactylus_benguellensis
Hemidactylus_ansorgii
Hemidactylus_greefii
Hemidactylus_brasilianus
Hemidactylus_palaichthus
Hemidactylus_agrius
Hemidactylus_laevis
Hemidactylus_ophiolepoides
Hemidactylus_isolepis
Hemidactylus_funaiolii
Hemidactylus_squamulatus
Hemidactylus_modestus
Hemidactylus_barbierii
Hemidactylus_lanzai
Hemidactylus_ophiolepis
Hemidactylus_albopunctatus
Hemidactylus_somalicus
Hemidactylus_citernii
Hemidactylus_foudaii
Hemidactylus_sinaitus
Hemidactylus_afarensis
Hemidactylus_pumilio
Hemidactylus_inintellectus
Hemidactylus_granti
Hemidactylus_dracaenacolus
Hemidactylus_luqueorum
Hemidactylus_persicus
Hemidactylus_hajarensis
Hemidactylus_yerburii
Hemidactylus_montanus
Hemidactylus_jumailiae
Hemidactylus_macropholis
Hemidactylus_pauciporosus
Hemidactylus_granchii
Hemidactylus_barodanus
Hemidactylus_mindiae
Hemidactylus_lemurinus
Hemidactylus_turcicus
Hemidactylus_lavadeserticus
Hemidactylus_dawudazraqi
Hemidactylus_paucituberculatus
Hemidactylus_masirahensis
Hemidactylus_inexpectatus
Hemidactylus_minutus
Hemidactylus_homoeolepis
Hemidactylus_oxyrhinus
Hemidactylus_forbesii
Hemidactylus_alkiyumii
Hemidactylus_shihraensis
Hemidactylus_festivus
Hemidactylus_saba
Hemidactylus_granosus
Hemidactylus_asirensis
Hemidactylus_alfarraji
Hemidactylus_ulii
Hemidactylus_almakhwah
Hemidactylus_awashensis
Hemidactylus_robustus
Hemidactylus_farasani
Hemidactylus_mandebensis
Hemidactylus_adensis
Hemidactylus_platyurus
Hemidactylus_craspedotus
Hemidactylus_bowringii
Hemidactylus_karenorum
Hemidactylus_garnotii
Hemidactylus_aquilonius
Hemidactylus_gujaratensis
Hemidactylus_giganteus
Hemidactylus_yajurvedi
Hemidactylus_hemchandrai
Hemidactylus_leschenaultii
Hemidactylus_flaviviridis
Hemidactylus_frenatus
Hemidactylus_treutleri
Hemidactylus_varadgirii
Hemidactylus_tenkatei
Hemidactylus_murrayi
Hemidactylus_chipkali
Hemidactylus_parvimaculatus
Hemidactylus_kushmorensis
Hemidactylus_gleadowi
Hemidactylus_xericolus
Hemidactylus_flavicaudus
Hemidactylus_sataraensis
Hemidactylus_vijayraghavani
Hemidactylus_reticulatus
Hemidactylus_gracilis
Hemidactylus_albofasciatus
Hemidactylus_prashadi
Hemidactylus_sushilduttai
Hemidactylus_kangerensis
Hemidactylus_maculatus
Hemidactylus_sahgali
Hemidactylus_whitakeri
Hemidactylus_triedrus
Hemidactylus_lankae
Hemidactylus_siva
Hemidactylus_hunae
Hemidactylus_scabriceps
Hemidactylus_depressus
Hemidactylus_kolliensis
Hemidactylus_pakkamalaiensis
Hemidactylus_graniticolus
Hemidactylus_vanam
Hemidactylus_paaragowli
Hemidactylus_easai
Hemidactylus_hegdei
Hemidactylus_acanthopholis
Hemidactylus_tamhiniensis
Hemidactylus_aaronbaueri
40
30
20
10
0
Millions of years ago

**DIVALIKE+J dispdistadj**
**ancstates: global optim, 2 areas max. d=0.0035; e=0; j=0.0044; LnL=-69.13**

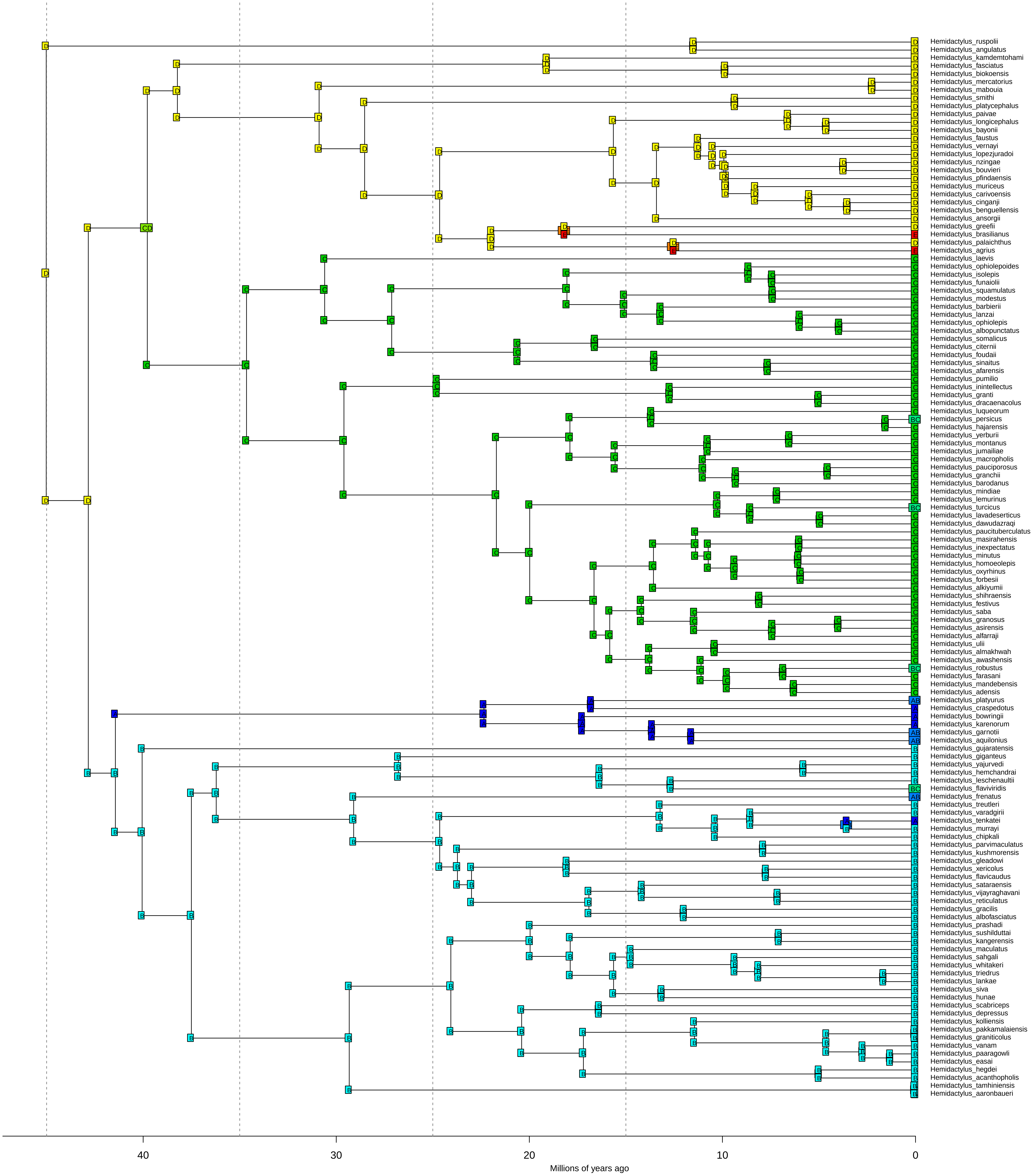

**DIVALIKE+J dispdistadj**

**ancstates: global optim, 2 areas max. d=0.0035; e=0; j=0.0044; LnL=−69.13**

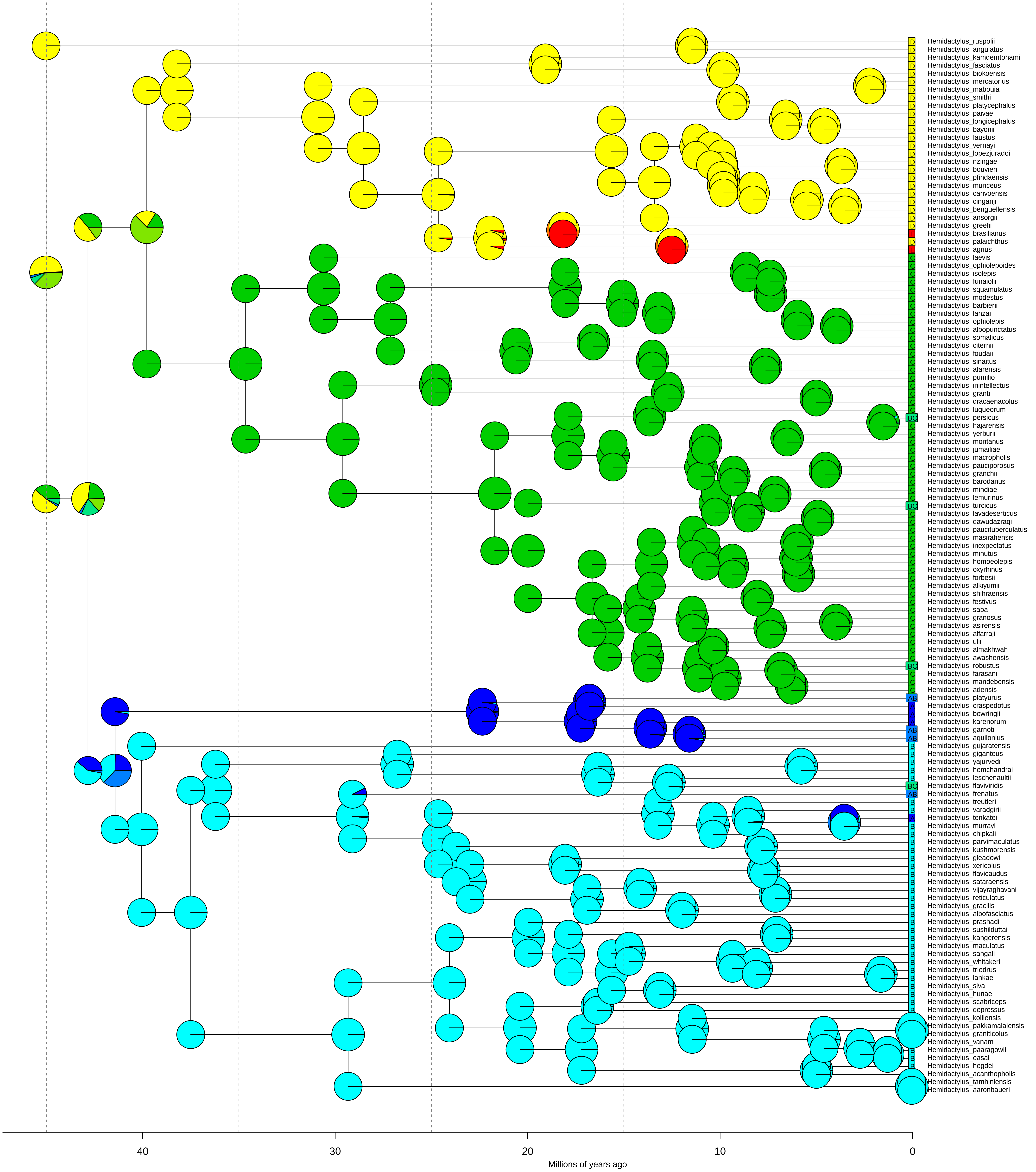
